# Endogenous retroviral elements LTR8B and MER65 regulate the PSG9 locus that promotes trophoblast syncytialization: Insights into placental evolution and pre-eclampsia pathology

**DOI:** 10.64898/2026.01.20.700661

**Authors:** Manvendra Singh, Yuliang Qu, Amit Pande, Julianna Zadora, Florian Herse, Martin Gauster, Xuhui Kong, Rongyan Zheng, Rabia Anwar, Katarina Stevanovic, Ralf Dechend, Marie Cohen, Attila Molvarec, Jichang Wang, Miriam K. Konkel, Bin Zhang, Cedric Feschotte, Gabriela Dveksler, Sandra M. Blois, Laurence D. Hurst, Zsuzsanna Izsvák

## Abstract

**Background:** Understanding the causes of the exceptional rate of evolution of the mammalian placenta is likely to aid the understanding of placental development and the aetiology of the human-specific pregnancy disorder pre-eclampsia (PE). As retroelements (REs) are often lineage-specific and known to be co-opted for placental functioning, here we consider the RE binding of GATA3 and DLX5, these transcription factors being dysregulated in PE, and their downstream consequences.

**Methods:** Multiomics analyses identified the retroviral regulatory sequence LTR8B in the PSG gene array, as a contributor to expression diversification in the placenta. To characterize this genomic domain, we performed copy number variation analysis and whole-genome sequencing. Multiomics data was employed to identify loci that might act as an active chromatin loop boundary around the PSG region. CRISPR-Cas9 knockouts with aligned RNAseq and epigenetic mark data tested for trophoblast-specific cis-regulatory elements (CREs-enhancer and/or promoter sequences) of resulting loci. Functional assays were employed to characterize the phenotypic effects of a candidate locus. Structural analysis of PSG family members also identified an additional RE, MER65-int. RNA-seq and antibody staining was employed to consider polyadenylation and functional diversification.

**Results:** The LTR8B CRE facilitates the binding of transcription factors (e.g., GATA3, DLX5, TFAP2A/C), resulting in a diversified PSG gene expression pattern within a primate-specific genomic region that exhibits high intraspecies variability. The LTR8B/PSG9 regulatory element influences other PSG family members. PSG9, unique among PSGs, produces both secreted and membrane-anchored isoforms, MER65-int providing alternative polyA signals, enabling the evolution of secreted PSG variants through the truncation of the ancestral CEACAM protein’s transmembrane domain. The LTR8B/PSG9 locus regulates the differentiation of multinucleated trophoblasts (syncytialization) and, like chorionic gonadotropin and syncytin1, determines the identity of syncytiotrophoblasts. Notably, PSG9 is the most upregulated PSG in PE, with levels correlated with GATA3 and DLX5 levels.

**Conclusions:** REs contribute to the structural and expression evolution of PSG genes, facilitating lineage-specific placental evolution. The LTR8B/PSG9 regulatory network plays a central role in syncytiotrophoblast differentiation. Given the association between DLX5/GATA3 dysregulation and elevated PSG9 levels, along with PSG9’s expression in the first trimester, PSG9 shows potential as a predictive biomarker for PE.

## Background

Of all organs, the placenta is exceptional in being morphologically fast-evolving with species-specific features [1–7]. Understanding the mechanistic basis of such rapid evolution should thus provide an exemplar of rapid organ-level evolution. Classically, we consider that adaptive evolution features are either changes to genes or changes in their expression [8]. But if expression changes over time, how might this occur? If proteins (or transcripts more regularly) change in length, for example, how might this occur?

In addition to this rapid evolution, dysfunction of the human placenta underpins the human-specific disease of pregnancy, pre-eclampsia (PE), defined as the manifestation of new-onset hypertension along with proteinuria or other maternal end-organ damages [9]. Affecting 3-8% of pregnancies worldwide, PE is a severe, heterogeneous, poorly understood condition [10–13]. PE is classified into early onset (EO-PE) or late onset (LO-PE) [14, 15], the former being especially dangerous to mother and infant [16–18].

A major challenge in the understanding of PE is that it has only been observed in humans [19]. While improved understanding of PE is important for both improved diagnostics and therapies, how to study a human-specific condition remains challenging, not least because of the lack of a robust animal model. One possible method of analysis is to use human-specificity as a strength rather than a weakness: we can incorporate an evolutionary dimension to identify lineage-specific pathways that are strong candidates for a lineage-specific disorder. Here, we presume that we can expect to better understand PE by searching for the mechanisms of rapid organ-level evolution.

The distribution of transposable elements often varies between species and tends to be specific to particular evolutionary lineages. Their co-option has the potential to explain rapid evolution and lineage specific traits. Endogenous retroviruses (ERVs) have been observed to influence gene expression and cellular processes during placental development. Certain ERVs have been repurposed and now play a role in the placentation process, so that their influence is directly linked to the evolutionary adaptation of the placenta [20–30]. These co-option events contribute to the species-specific characteristics of placentation (e.g. depth of trophoblast invasion, regulation of immune responses and vascular function, etc.) [31]. For example, ERV-derived sequences (e.g., long terminal repeats (LTRs)) serve as regulatory elements for gene expression by providing enhancers and signals for splicing and polyadenylation [27, 30, 32–35]. The co-option of other retroviral sequences, such as open reading frames (ORFs), may also lead to novel placental genes, such as the retroviral envelope-derived placental syncytins and suppressyn [20–23, 28, 36, 37]. Since ERVs have been shown to influence important processes in the human placenta, we hypothesise that their dysregulation may contribute to the development of pathological conditions such as PE.

We start by asking which ERVs are bound by placental transcription factors (TFs), including our two prior top candidates for human TF genes that are overexpressed in PE, GATA3 and DLX5 [38]. The expression of both transcription factors marks preimplantation trophectoderm (TE) and likely contributes to the regulatory network governing the placenta development [39–43]. While GATA3 regulates the differentiation of trophoblast progenitors in both mice and humans [44], DLX5 is not expressed in TEs of rodents, consistent with the possibility that regulatory networks (regulons) involving these genes are evolutionarily younger. We identify several ERV families that have significant co-occupancy with these TFs.

Might the dysregulation of both TFs have consequences for the function of ERVs that remodel the placental gene regulatory network of PE? To address this, we investigated the overlap between the GATA3 / DLX5 target genes, ERVs and clinical data from PE patients. This approach led us to the recently expanded gene array encoding pregnancy-specific glycoproteins (PSGs) [45] on human chromosome 19q13.

PSGs are *a priori* likely players in lineage-specific effects as they are unusually fast evolving in many dimensions (e.g. copy number, domain structure) [45]. Indeed, the PSG locus is among the fastest-evolving genomic regions in the human genome [46, 47]. PSGs and the carcinoembryonic antigen-related cell adhesion molecules (CEACAMs) belong to the carcinoembryonic antigens (CEA), members of the immunoglobulin (Ig) superfamily [48, 49]. While the ancestral gene of CEACAM/PSG is thought to be common to both primates and rodents, it was subject to convergent evolution [48], accompanied by a series of gene duplications [50] and subsequent diversification events, resulting in CEACAM and PSG gene clusters. The CEACAM proteins are cell membrane-anchored proteins, whereas PSGs are commonly secreted into the maternal circulation [48, 49]. The PSG gene clusters are expanded independently in mouse and human, represented by Psg1-17 and PSG1-PSG11, in mouse and human, respectively [51–54]. Thus, the human PSG genes have no orthologs in non-primates [45, 55].

PSGs are also likely to be involved in placental evolution and placental disorders given what we know of their biology. PSGs are, for example, implicated in key trophoblast functions, including the establishment of maternal immune tolerance to the foetus [56–58]. While PSGs are expressed in both extravillous trophoblast (EVTB) and syncytiotrophoblast (STB) [12, 59], many of their characterized functions (e.g. regulating angiogenesis, cell adhesion and migration [43, 58–63]) are typical of the EVTB, whereas their role(s) in STBs are less well understood. Not all but certain family members have been associated with pathological pregnancies. Evidence concerning their contribution is however complex and contradictory [45, 59, 64]. For example, reduced PSG1 and PSG11 expression [65] and elevated levels of PSG7 and PSG9 are suggested to confer the risk for PE [66–68].

Here then we consider the role of HERVs in the evolution and activity of PSGs. We find that a combination of HERVs (e.g. LTR8B and MER65-int (MER65 thereafter)) have been co-opted at the PSG locus. Our integrative analysis of single-cell transcriptome and epigenome data, confirmed by experimental validation, indicates that PSGs are regulated by trophoblast-specific transcription factors (GATA3, DLX5, TFAP2A/C) whose binding sites are embedded in the LTR8B sequence, which we recognised as trophoblast-specific candidate (c)CRE. MER65 provides alternative polyA signal(s), thus contributing to the evolution of novel protein variants, secreted into the maternal circulation.

We find that the PSG9 locus controlled by the cis-regulatory element (CRE), LTR8B, appears to be exceptional within the PSG array acting as a master regulator of the array. CRISPR-Cas9-mediated knockouts, and 3D chromatin studies show that LTR8B/PSG9 serves as a binding platform for a specific set of placenta-specific TFs and defines the 3D domain boundary. While acting as a strong CRE for PSG9 expression, it regulates the differentiation programme of human multinucleated trophoblasts (syncytialization). Furthermore, in contrast to the other family members that are exclusively secreted, PSG9 possesses both secreted and membrane-anchored isoforms. By comparing multiomic data with transcriptomes collected from PE patients, we establish a link between dysregulated DLX5/GATA3 and elevated levels of the secreted PSG9 protein. This holds the potential of a new predictive biomarker for the human-specific pregnancy disease pre-eclampsia.

## Methods

### Cell culture and transfection

BeWo cells were maintained in GlutaMAX™ DMEM/F-12 medium (#31331028, Thermo Fisher Scientific) supplemented with 10% (v/v) fetal bovine serum (FBS) and 1% (v/v) penicillin/streptomycin (P/S). Saint George’s Hospital Placental cell Line-4 (SGHPL-4) cells were cultured in Ham’s F-10 liquid medium (#FG0715) supplemented with 10% FBS and 1% P/S. All cells were cultured in a humidified incubator at 37°C with 5% CO₂. Transfections were performed using Lipofectamine™ 3000 reagent (#L3000015, Thermo Fisher Scientific) for BeWo cells and the Neon™ Transfection System (#MPK5000, Thermo Fisher Scientific) for SGHPL-4 cells, following the manufacturers’ instructions. To generate stable cell lines using the Sleeping Beauty (SB) system [69], transfected cells were further selected in complete medium containing approximately 4 µg/ml puromycin for 10–14 days.

The human trophoblast stem cells (hTSCs) were derived from our previous study [70]. Briefly, hTSCs were seeded onto 1% Matrigel–coated surfaces and cultured in hTSC medium [71] with minor modifications. The medium consisted of DMEM/F12 (#11320033, Thermo Fisher Scientific) supplemented with 0.2% FBS (#FSP500, ExCell Bio), 1% ITS-X (#51500056, Thermo Fisher Scientific), 0.1 mM 2-mercaptoethanol (#21985023, Thermo Fisher Scientific), 0.3% BSA (Thermo Fisher Scientific, Cat#15260037), 50 µg/ml L-ascorbic acid (Sigma, Cat#A4544), 50 ng/ml human EGF (#AF-100-15, PeproTech), 2 µM CHIR99021 (#S1263, Selleckchem), 0.5 µM A83-01 (#2939, Tocris), 1 µM SB431542 (#S1067, Selleckchem), 10 µM VPA (#S3944, Selleckchem), 5 µM Y27632 (#S1049, Selleckchem), and 100 µg/ml Primocin (#Ant-pm-2, InvivoGen). Cells were routinely maintained at 37 °C in 5% CO₂, with medium changes performed every other day.

The STB induction from hTSC was performed following the published protocol [71]. The hTSC cells were seeded on 1% Matrigel-coated 12-well plate (5×10^4^ cells per well) and cultured in the STB differentiation medium: DMEM/F-12 supplemented with 0.1 mM 2-mercaptoethanol, 100 μg/ml Primocin, 0.3% BSA, 1% ITS-X supplement, 2.5 μM Y27632, 2 μM forskolin, and 4% KnockOut Serum Replacement (KSR). The STB medium was changed daily.

### ChIP-exonuclease (exo) assay

Generation of a stable TET-inducible SGHPL-4 cell line expressing DLX5: SGHPL-4 cells were transfected with the doxycycline-inducible vector pTOVT11-HA-DLX5-(SV40-puro). A total of 1 × 10⁶ SGHPL-4 cells (passage 13) were electroporated using the Neon Transfection System (1260 V / 20 ms / 2 pulses) with 3 µg of the expression vector and 300 ng of the *Sleeping Beauty* transposase plasmid (pcGlobin2-SB100X). Two days post-transfection, cells were subjected to puromycin selection (3 µg/ml) for 12 days. Stable cells were frozen at passage 18/19. For the ChIP-exo assay, 2 × 1.5 × 10⁶ SGHPL-4 pTOVT11-HA-DLX5-(SV40-puro) cells were seeded into two 15 cm dishes in 15 ml of Ham’s F-10 medium supplemented with 10% fetal calf serum (FCS) and amino acids (AA), and incubated overnight at 37°C in 5% CO₂. When cells reached ∼70–75% confluency, the medium was replaced with fresh Ham’s F-10 medium (+10% FCS, +AA). DLX5 expression was induced by adding doxycycline (2 mg/ml stock; 5 µl per 10 ml of medium). The ChIP-exonuclease assay was performed according to the method described by Serandour et al. [67, 72]. Libraries were quantified using the KAPA Library Quantification Kit for Illumina platforms (KAPA Biosystems, KK4824) and sequenced on a HiSeq system following the manufacturer’s protocol.

### Transient and stable overexpression of gene of interest (GOI)

For transient overexpression, the gene of interest (GOI) was cloned into the *Sleeping Beauty* (SB) pT2 vector (5′-IR-CAGGS promoter-HA-MCS-SV40 promoter-Puro-3′-IR). The resulting plasmids were introduced into target cells via transfection. At 24 hours post-transfection, cells were selected in complete medium containing approximately 4 µg/ml puromycin for 48 hours. Transfected cells were then harvested for downstream applications, and GOI overexpression was confirmed by RT-qPCR or western blotting. Cells transfected with a plasmid encoding the mCherry fluorescent protein, prepared using the same protocol, served as a negative control. To generate stable cell lines expressing the GOI, cells were co-transfected with the GOI expression construct and the SB transposase plasmid (pcGlobin2-SB100X) at a 10:1 ratio. The experimental procedure was otherwise identical to that used for transient overexpression. For stable overexpression, 2.5 µg of the GOI expression cassette and 250 ng of pcGlobin2-SB100X were co-transfected per well (note: double-check if “250 µg” in the original was a typo; if not, adjust accordingly). Stable integration was selected with puromycin, and successful overexpression was validated as described above.

### Quantitative RT-qPCR

The RNA extraction from cells was carried out with the Direct-zol™ RNA MiniPrep Kit (#R2025 Zymo Research). Then 500 ng ∼ 1 µg total RNA was used to synthesize single-stranded cDNA with the High-Capacity RNA-to-cDNA™ Kit (#4387406 Thermo Fisher Scientific) following the manufacturer’s instructions. The resulting cDNA was used for real-time qPCR with the SsoAdvanced™ universal SYBR^®^ Green Supermix (#1725271 Bio-Rad) on the CFX96 Touch^TM^ system (Bio-Rad). Gene expression was analysed by the CFX Maestro^TM^ Software (Bio-Rad) with the ΔΔCT method and normalized to human *ACTIN* expression. The RT-qPCR with human placenta samples was done following the protocol in the previous study. For the primers, see Additional File 1.

### Western blotting

Cell lysis was performed at 4°C for 2 hours using a lysis buffer containing 50 mM Tris-HCl (pH 8.0), 100 mM NaCl, 5% glycerol, 10 mM EDTA, 1% NP-40, and an appropriate amount of protease inhibitors (#A32955, Thermo Fisher Scientific). The lysates were then centrifuged at 14,000 rpm for 10 minutes at 4°C, and the supernatants were collected for downstream applications. Protein concentrations were determined using the Pierce™ BCA™ Protein Assay Kit (#23225, Thermo Fisher Scientific). Protein samples were denatured by boiling in SDS sample buffer at 96°C for 10 minutes. SDS-PAGE was performed using the TGX Stain-Free™ FastCast™ acrylamide kit (#1610183, Bio-Rad), following the manufacturer’s instructions. Proteins were transferred to membranes using the Trans-Blot® Turbo™ Transfer System (Bio-Rad). Membranes were blocked in 5% (w/v) skim milk in TBST for 1 hour at room temperature (RT), followed by incubation with the primary antibody overnight at 4°C. After washing the membranes with TBST (6 times, 5 minutes each), they were incubated with the secondary antibody for 1 hour at RT. The membranes were then washed again (6 × 5 minutes) in TBST. Chemiluminescent signals were developed using the ECL™ detection reagent (#RPN2232, GE Healthcare) and visualized using the Bio-Rad ChemiDoc™ MP Imaging System. Band intensities were quantified by densitometric analysis using Image Lab™ software (Bio-Rad), according to the manufacturer’s instructions.

### Checking the specificity of the PSG9 antibody

The specificity of the PSG9 antibodies were tested (Additional File 2: Fig. S2): (Left panel) Purified recombinant PSG proteins (1.5 μg / lane) (PSG1 V5His, PSG2 V5His, PSG4 V5His, PSG5 V5His, PSG6 V5His, PSG7 V5His, PSG8 V5His, PSG9 V5His and PSG11) [62] were loaded on the SDS-PAGE gel. Following protein transfer the membrane was first blocked in a solution (2.5% milk 0.1% in TBS-T) 1h @RT. The Abs were: Ab Novus rabbit polyclonal NBP-2 19979 at a 1:1,000 dilution, followed by goat anti-rabbit HRP conjugated Ab. Note that this Ab is advertised as PSG9-specific but it is not [45]. (Right panel) 1.5 μg / lane of purified recombinant PSG proteins PSG1 V5His, PSG2 V5His, PSG4 V5His, PSG5 V5His, PSG6 V5His, PSG7 V5His, PSG8 V5 His, PSG9 V5His and 20 μg lysates of BeWo, SHGPL-4 cells were loaded on 12% SDS-PAGE gel. Following protein transfer the membrane was first blocked in a solution (2.5% milk 0.1% in TBS-T) 1h @RT, than incubated with the rabbit polyclonal anti-PSG9 (ab64425, Abcam) 1:1000 in 2.5% milk (0.1% in TBS-T) incubation, O/N 4°C, followed by anti-rabbit-HRP 1:5000 in 2.5% milk (0.1% in TBS-T) incubation: 1h @RT. Specificity: No cross-reactivity was observed with other PSGs. Bands are detectable only in the lane loaded by recombinant PSG9 and in the BeWo and SGHPL-4 cell lysates.

### Sandwich ELISA to determine circulating PSG9 levels

PSG9 concentrations in the serum of pregnant woman were determined by a specific sandwich ELISA. The NBP1-57676 capture Ab recognizes the synthetic peptides corresponding to PSG9 (pregnancy specific beta-1-glycoprotein 9). The peptide sequence YSNASLLIQNVTRKDAGTYTLHIIKRGDETREEIRHFTFTLYLETPKPYI was selected from the N terminal of PSG9. We used a N-domain PSG9 mutant as internal control in our ELISA. Briefly, immunolon 2 ELISA plates (Dynatech Laboratories, USA) were coated with anti-PSG9 antibody (5 μg/ml; (NBP1-57656, Bio-Techne, Germany) and washed with washing buffer (0.5% Tween-20 in PBS). Plates were blocked with 3% BSA in PBS. Individual wells were incubated with serial dilutions of PSG9-Fc [73] or serum samples (diluted 1/10) for 2 h at room temperature (RT). Wells were washed and incubated with HRP-conjugated anti-PSG mAb-BAP3 [45], which recognizes the B2 domain of all human PSGs (0.25 μg/ml in PBS 0.1% BSA; sc-59348 Santa Cruz Biotechnologies, USA). After eight additional washes, a colorimetric reaction was developed with the 3,3,5,5′-tetramethyl benzidine (TMB) substrate (Pierce Biotechnology, USA). The reaction was stopped by adding one volume of 4 N H_2_SO_4_. Absorbance at 450 nm was recorded. Each reported value is the mean of triplicate assays. Specificity: The NBP1-57676 did not recognizes the N-domain PSG9 mutant. No cross-reactivity was observed with a mutant of PSG9 lacking the N-domain (Dveksler, unpublished), PSG1, PSG2, PSG4, PSG5, PSG6, PSG7, PSG8, PSG11, CECAM1, and CECAM5) [62].

### Cell fractionation assay

To examine the subcellular localization of PSG9 isoforms, BeWo cell lines were generated to stably co-express two representative PSG9 isoforms, each fused to either a FLAG or HA epitope tag. Transfected cells were seeded in 6-well plates at a density of 1 × 10⁶ cells per well and cultured for 48 hours. Approximately 4 ml of culture medium was collected from each well and centrifuged at 4,000 rpm for 15 minutes at 4°C. The resulting supernatant was concentrated using the Amicon® Ultra-15 Centrifugal Filter Unit (#UFC903008, Merck) for subsequent analysis. For cellular compartment fractionation, approximately 1 × 10⁷ cells were harvested and processed using the Minute™ Plasma Membrane Protein Isolation and Cell Fractionation Kit (#SM-005, Invent Biotechnologies), following the manufacturer’s instructions. The resulting fractions—culture medium, total cell lysate, plasma membrane, and cytoplasmic components—were analyzed by Western blotting. Membranes were incubated overnight at 4°C with a primary antibody mixture containing anti-FLAG and anti-HA antibodies. After washing, membranes were incubated with the appropriate secondary antibody mixture, and chemiluminescent signals were developed for detection.

### Knocking down of PSG9 expression

BeWo Cells were transfected with the plasmid pLKO.1-shRNAs (PSG9 MISSION^®^ shRNA TRCN0000244632, CGAGGTGATGAGACTAGAGAA). The 21nt oligonucleotide has 100% identity to PSG9 (score 42.1 bits(21)). The next potential target was PSG3 with the score 30.2 bits(15). The pLKO.1-scramble shRNA (#1864 Addgene).

### RNA-seq of trophoblast from placenta

Total RNA was isolated from tissues (homogenized by ceramic beads) and cells using QIAzol lysis reagent and Qiagen RNeasy mini kit (including on-column DNAase I digestion) (Qiagen) according to the manufacturer’s protocol. RNA quality and concentration was measured by NanoDrop-1000 spectrophotometer (PeqLab). RNA was reverse transcribed into cDNA using High-Capacity cDNA Reverse Transcription Kit (Applied Biosystems) and detected by real-time polymerase chain reaction (PCR) on ABI 7900HT Fast Sequence Detection System (Applied Biosystems). Data was analysed by 7900HT Fast System Software (Applied Biosystems). Primers were designed with Primer3Plus, GeneScript Real-time PCR Primer Design tool, PrimerExpress 3.0 (Applied Biosystems) or we used already published primers. Primers were synthetized by Biotez (Germany). The expression of all of the genes was normalized to 18S expression.

### Immunohistochemistry

Human FFPE placental tissue sections from first trimester and term pregnancies were mounted on Superfrost Plus slides. Sections underwent standard deparaffinization, followed by immunostaining with the UltraVision Large Volume Detection System HRP Polymer Kit, according to the manufacturer’s protocol. Briefly, endogenous peroxidase was blocked with UltraVision hydrogen peroxide block for 10 min. Slides were washed three times with TBS containing 0.05% Tween 20 (TBS/T). This was followed by incubation with Ultra Vision Protein Block for 5 minutes. Polyclonal anti-PSG9 antibody (8µg/ml, ab64425, Abcam) was diluted in Antibody Diluent (DAKO) and applied to the slides, which were incubated for 45 minutes at room temperature (RT). After three washing steps with TBS/T, detection was performed using a primary antibody enhancer and an HRP-labeled polymer system, incubated for 15 minutes. This was followed by development with 3-amino-9-ethylcarbazole (AEC, Thermo Scientific) according to the manufacturer’s instructions. Nuclei were counterstained with hematoxylin, and slides were mounted using Kaiser’s glycerol gelatin (Merck). For negative controls, slides were incubated with Negative Control for Rabbit IgG Ab-1 (NeoMarkers/Thermo Scientific) at the same concentration as mentioned above. Images were acquired with a Leica microscope (Leica DM6000B) and a digital camera (Olympus DP72).

### Immunofluorescence staining

Cells were seeded onto coverslips into 12 well plate (4 × 10^5^ per well) for 48 hr, fixed with 4% paraformaldehyde in PBS for 15 min at RT, permeabilized with 0.2% triton X-100 in PBS for 15 min at RT (optional for intracellular staining), and blocked with 10% serum solution. The coverslips were incubated with the primary antibody for overnight at 4 °C and washed three times with PBS. Then the second antibody solution was added. The Nuclear staining was done with1 µg/ml Hoechst 33342 (#H3570 Thermo Fisher Scientific) for 5 min at RT and the coverslips was mounted with ∼ 15 µl mounting medium (#H-1000 Vector Laboratories). Images were captured under the Leica TCS SP8 with a 63x oil/NA 1.4 objective at the Advanced Light Microscopy (ALM) technology platform at the Max Delbrück Center for Molecular Medicine (MDC).

### Luciferase reporter assays

To characterize the *upstream regulatory* TEs elements associated with PSGs expression, reporter assay containing various RE sequences upstream of the luciferase gene was designed. The candidate RE sequences were introduced into the pGL3 Promoter Vector MCS-SV40*^prom^*-luc-SV40pA (#E1761 Promega). For transient transfection, BeWo cells, seeded onto 96-well plates (4 × 10^4^ cells/well) for 24 hr were co-transfected with the internal control pRL-SV40 Vector (50 ng/well, #E2231 Promega) and candidate plasmids (250 ng/well) using the jetPRIME^®^transfection reagent (#114-15, Polyplus Transfection) following the manufacturer’s instructions. At 48 hr post-transfection, transfected cells were harvested and lysed. The resultant lysate was used to measure the *firefly* (luc) and *Renilla* (Rluc) luminescence with the Dual-Glo^®^ Luciferase Assay System (#E2920 Promega) on the Tecan Spark 10M multimode plate reader. The average luc/Rluc luminescence ratio of at least 6 technical wells was calculated to represent the relative enhancer activity of candidate elements. The empty vector and the pGL3-Control Vector containing an SV40 enhancer (#E1741 Promega) were used as the negative and positive controls, respectively.

For the *polyadenylation* reporter assay, candidate elements were cloned into the dual luciferase vector SV40*^prom^*-Rluc-MCS-IRES-luc-SV40pA (kindly offered by Prof. Dr. Xianchun Li) [74]. For transient transfection, BeWo cells seeded onto 96-well plates (4 × 10^4^ cells/well) for 24 hr were transfected with the control or candidate plasmids (400 ng/well) using the jetPRIME^®^ transfection reagent. At 48 hr post-transfection, transfected cells were harvested and lysed. The luminescence signals were developed following the protocol aforementioned. The average luc/Rluc luminescence ratio of at least 7 technical wells was calculated to represent the relative polyadenylation efficiency of candidate elements. The empty vector and the one containing a known synthetic pA site (SPA) were used as the negative and positive controls, respectively.

### Characterization of the chromatin features of the human PSG gene array

ChIP-seq datasets representing transcription factors (TFs), histone modifications, and regulatory complexes in the differentiated and isolated trophoblast cells were retrieved from GEO GSE databases (referenced in the main text). ChIP-seq reads were aligned to the Hg19 human reference genome using the Bowtie2. All reads with a phred score less than 33 and PCR duplicates were removed using bowtie2 and Picard tools, respectively. ChIP-seq peaks were called by MACS2 with the parameters in “narrow” mode for TFs and “broad” mode for histone modifications, keeping FDR < 1%. ENCODE-defined blacklisted regions were excluded from called peaks. We then intersected these peaks with the loci from TE subgroups using ‘bedtools’ with any overlap. For ChIP-seq binding enrichment on a subset of marks following motif analysis, 70% overlap of peak and TE was required. Enrichment of a given TF and histone marks within TE subgroups was calculated using ‘enrichR’ package in R, using the customized in-house codes (see the codes on GitHub for the detailed analysis pipelines and calculation of enrichment score). Note-Publicly available datasets were analysed using their originally aligned genome assemblies (hg19 or hg38), as specified in the respective repositories. Newly generated datasets were processed with hg38. Coordinate conversion between assemblies was intentionally avoided to prevent artefacts in repetitive regions, particularly over transposable elements.

### Identification of polyadenylation *cis*-element in MER65

Each MER65 sequence fused to the 3’-UTR of the PSG transcript was retrieved as a query sequence (200 bp long, centred at the polyadenylation site). Then the query sequence was used to identify its corresponding alignment in the MER65 consensus sequence. The identification of polyadenylation *cis*-element was performed following the protocol in [75].

### Analysis of the distribution pattern of MER65 elements across the human genome

All MER65 elements were extracted from masked (RepeatMasker version 4.0.5 repeat Library 20140131) GRCh38/hg38 (alt chromosomes removed). Paralelly, the coordinates of genes in gene track format (gtf) were downloaded from hg38 RefSeq databases. All annotated MER65 elements were intersected with the gene start and end sites coordinates using ‘intersectbed’ and the distance of MER65 was measured using the ‘closestBed’ function inbuilt into the software ‘bedtools’.

### Primary human trophoblast isolation

Placental tissues from healthy and pre-eclamptic (PE) patients (n=5 each) were collected at HELIOS Klinikum, Berlin, following approved protocols (Charité Berlin). All placentas were processed within 2 hours post-delivery according to [76]. Cell purity (>92%) was confirmed by cytokeratin-7 staining and flow cytometry. Cells were incubated overnight at 37°C, then collected and stored at –80°C for RNA, DNA, and protein isolation.

### Primary human trophoblasts mRNA isolation, Library preparation and RNA-sequencing

Total RNA was extracted from primary human trophoblasts (8 healthy and 10 early-onset PE samples) using TRIzol reagent and purified with the Direct-zol™ RNA MiniPrep kit (Zymo Research), including on-column DNase I digestion. RNA concentration was measured using a NanoDrop ND-1000, and quality assessed with the Agilent 2100 Bioanalyzer and RNA 6000 Nano Kit. Libraries were prepared using the Illumina TruSeq Stranded mRNA LT Set A kit (550 ng input RNA/sample), with unique sample indices allowing for pooling. Sequencing was performed as 100 bp strand-specific paired-end reads on the Illumina HiSeq 2000 platform (BIMSB Genomics Platform, MDC Berlin). Clustering used the PE TruSeq Cluster Kit v3 on a cBot System. Demultiplexing and conversion to FASTQ files were done via CASAVA 1.8.2.

### Construction of the KO LTR8B cell line

Two pairs of sgRNAs flanking the LTR8B copy at the *PSG9* gene locus were designed with the online tool CRISPOR (http://crispor.tefor.net/) (Additional File 1) and cloned into the pU6-(BbsI) sgRNA_CAG-Cas9-venus-bpA (#86986 Addgene, a gift from Ralf Kuehn). Then the 5’ sgRNA and 3’ sgRNA plasmids were co-transfected into BeWo cells in 6-well plates (1.25 µg/well each). At 48 hr post-transfection, the Venus-positive cells were sorted into 6-well plates (∼ 5000 cells/well). After 14 days, single colonies were manually picked under microscope and genotyped using the DNA isolated with the QuickExtract™ DNA Extraction Solution (#QE09050 Lucigen). The resulting homozygous deletion colonies were further confirmed by sequencing and selected for the downstream usage. The wide type bulk cells were used as the negative control.

### Construction of the KD PSG9 cell line

The KD construct (21nt, PSG9 MISSION® shRNA TRCN0000244632, CGAGGTGATGAGACTAGAGAA) was specific to PSG9 (42.1 bits(21)) compared to the closest possible target (30.2 bits(15)). For the transcriptional analysis of KD PSG9 versus and KD scramble control, BeWo cells were transfected with the plasmid pLKO.1-PSG9 shRNA (MISSION^®^ shRNA TRCN0000244632). At 24 hr post-transfection, cells were screened with the 4 µg/mL puromycin for another 24 hr. Then the transfected cells were treated with the mixed medium containing 4 µg/mL puromycin and 50 µM Forskolin for 48 hr. Cell fusion measurement was done following the methods aforementioned. The pLKO.1-scramble shRNA (#1864 Addgene, a gift from David Sabatini) was used as a negative control.

### Fluorescence based BeWo cell syncytialization assay

For quantification of the syncytium, the BeWo cells stably expressing either the GFP or mCherry were co-seeded into 12-well plates with an equal number (1.5 × 10^5^ cells/cell type/well) for 24 hr. Then the culture medium was replaced with the fresh media containing 50 µM Forskolin (#F6886 Sigma-Aldrich). At 48 hr post-treatment, the co-cultured cells were harvested into 600 µl FACS buffer (DPBS containing 1% BSA and 5 mM EDTA) and filtered through the 70 μm Flowmi™ Cell Strainers (#15342931 Fisher Scientific) immediately before Fluorescence-Activated Cell Sorting (FACS) analysis. 20 ∼ 30 thousands of single cells were analysed for each sample and data analysis was done using the FlowJoTM software. The fusogenic capacity was evaluated based on the percentage of double positive cells.

### Transcriptomic profiling of KO LTR8B BeWo cells

To determine the transcriptomic profile of LTR8B knockout (KO) cells (sgRNA 392 + sgRNA 1073, P6-B4 colony) and a wild-type BeWo control, the cells were subjected to forskolin-induced synchronisation (see above). Total RNA was extracted from the cells 48 hours after forskolin treatment. Quality control (QC) of all RNA samples was performed using the Agilent 2100 Bioanalyzer System, including concentration, 28S/18S ratio and RNA integrity number (RIN). The transcriptome of the RNA samples was prepared using a stranded mRNA library kit and sequenced on the DNBseq platform at the BGI Group. Reads were aligned to the GRCh37/Hg19 reference genome using the STAR aligner [72]. The alignment quality was further assessed using the ‘RSeQC’ package [77].

### Analysis of RNA-Seq Data

Raw sequencing reads were processed using FASTX-Toolkit and Trimmomatic [78] to remove adaptor sequences, discard low-quality reads, and trim poor-quality bases. Outlier reads with more than 30% disagreement were excluded from further analysis. Transcript quantification was performed using Salmon [79]. Indexing was carried out with the command: salmon index -t transcripts.fa -i transcripts_index --decoys decoys.txt -k. Reads were aligned and quantified using: salmon quant -i transcripts_index/ −l IU −1 fastq −2 fastq --validateMappings -o output. Quality control checks for GC content and gene length bias were conducted using the NOISeq R package [80], which also generated quality diagnostic plots for count data. To assess replicate concordance, mean-variance analysis and Principal Component Analysis (PCA) were performed using the tximport package [81] in R. Analyses were based on lengthScaledTPM, with filtering thresholds of CPM >2 in at least two replicates per group. Batch effects were corrected using the RUv package [82] from Bioconductor. Normalization of expression data was then performed using the TMM (Trimmed Mean of M-values) method in preparation for differential expression analysis. Differential gene expression (DEG) was calculated using DESeq2 [83], employing default statistical settings. Additional statistical analyses were carried out in GraphPad Prism 9 (GraphPad Software, San Diego, CA, USA). Data are presented as mean ± SD, and statistical significance was defined as *P* < 0.05.

### Whole genome sequencing and analysis

Frozen placenta tissues were pulverized with liquid nitrogen. DNA from placentas was extracted using the DNeasy Blood and Tissue Kit (Qiagen) according to the manufacturer’s instructions. DNA was stored at −20°C. Genomic DNA samples were sequenced on an Illumina X HiSeq Platform. Each of samples were given a full lane on an 8 lane flowcell, running a paired end run with 150 bp reads. All sequenced samples met the initial DNA concentration thresholds and achieved >30x coverage of the Hg19 reference genome (specifics regarding kit/reagents/adapters can be provided by the sequencing facility at USUHS. All samples were aligned to the Hg19 human reference genome sequence using HAS (HiSeqAlignment Software) which comprises the Isaac aligner, Strelka SNP caller, and Manta/Canvas Structural variant caller. This pipeline is fairly robust and was able to accurately map reads across the PSG region based on the percent identify we observed between the copies of PSG across the locus. We also measured alignability across the locus. Lastly the samples had a high depth of coverage making accurate alignment possible. Specifically, this region was limited to chr19 40000000-48000000, which is 8 mb of sequence surrounding the PSG locus. The detected CNVs and SVs of 1000 bp or greater in size were split into PE and control groups.

### Trans-well invasion assay with SGHPL4 cells

The trans-well invasion assay using SGHPL4 cells was adapted from Angelova et al. (2013). Each trans-well insert was coated with 50 µl of 0.1 mg/ml Matrigel (diluted in chilled serum-free Ham’s F10 medium) and incubated at 37°C for ∼4 hours. For the chemoattractant, 600 µl of completed Ham’s F10 medium was added to the lower chamber. Human Epidermal Growth Factor (50 ng/ml) served as the positive control, while serum-free medium was used as the negative control.

SGHPL4 cells were trypsinized at 37°C for ∼3 minutes, resuspended in serum-free Ham’s F10 medium, and centrifuged. The cell concentration was adjusted to ∼1×10⁶ cells/ml, and 100 µl (∼1×10⁵ cells) were seeded in each insert. Plates were incubated at 37°C for 16 hours. For fixation, cells were washed in PBS, fixed with 3.7% PFA for 5 minutes, and permeabilized with 100% methanol for 20 minutes. After staining with 0.2% crystal violet for 30 minutes in the dark, non-invading cells were removed with a cotton swab. Invasive cells were visualized and counted in four microscopic fields (10x magnification).

## Results

### Identification of the ERV family LTR8B, co-occupied by DLX5 and GATA3, in placental gene regulation

To gain a comprehensive understanding of ERV-derived regulatory elements controlled by key trophoblast-specific transcription factors (TFs), we analysed ChIP-seq and ATAC-seq data from trophoblast stem cells (TSCs) and two differentiated cell types: TSC-derived extravillous trophoblasts (EVTBs) and trophoblast cells (TBs) differentiated from ESC_H1 (Fig. 1A). We began by identifying ERV families significantly enriched in genome-wide H3K4me1, H3K27ac, and H3K4me3 profiles in trophoblast cells to characterize their cCRE activity. We also analysed H3K9me3 signals, as most repressed ERVs are marked by this histone modification [84]. Additionally, we integrated ATAC-seq data from TSCs and their EVTB derivatives to identify genomic regions exhibiting accessible chromatin. Analysing ChIP-seq data for P300 and MED1, which are hallmarks of active enhancers [85] and ASCL2 and TEAD4, which play a pivotal role in trophoblast differentiation [86, 87] provided an integrated view of transcriptional regulatory elements (TREs), transcription factor (TF) binding, and gene regulation within the context of trophoblast development. When exploring trophoblast-specific features, we also examined TF data for TFAP2A, TFAP2C, DLX5 and GATA3 in trophoblast cells [84, 88] (Fig. 1B).

**Fig. 1.**
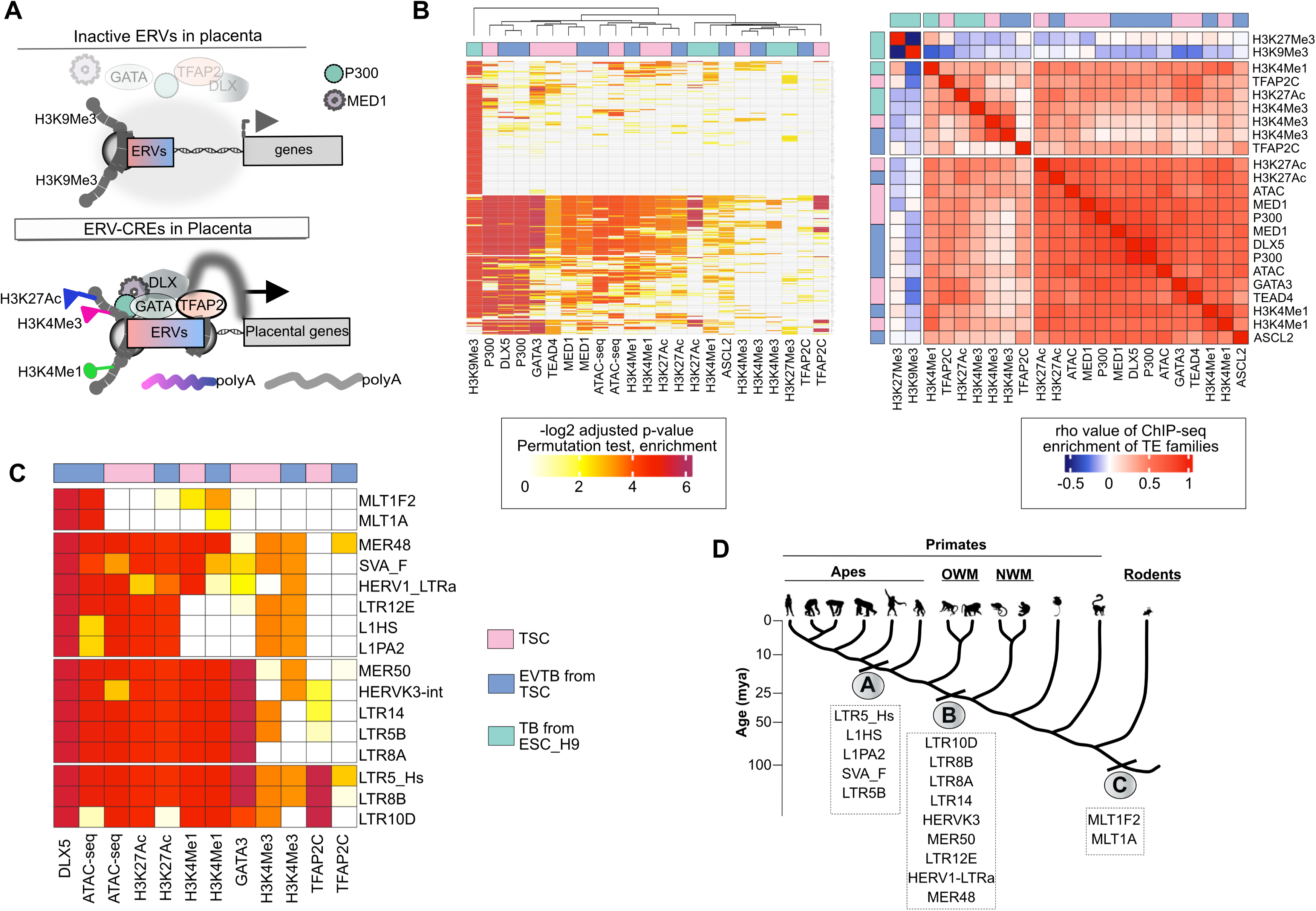
GATA3 and DLX5 are evolutionary new-comers in the human genome and are frequently associated with ERV-derived sequences. **A.** Schematic illustrating the interplay of histone modifications, transcription factor (TF) binding, accessible chromatin, and their regulators with endogenous retroviral (ERV) elements, which together govern the expression of nearby placenta-specific genes. This strategy is employed to catalog functional ERV-derived cis-regulatory elements (CREs) that regulate placental gene expression. **B.** Identification of ERVs significantly enriched in genome-wide binding profiles of the indicated TFs, histone modifications, and their regulators. Enrichment of each dataset within ERV subgroups was calculated using the enrichR package in R, with customized in-house scripts (see the GitHub repository for detailed analysis pipelines and enrichment score calculations). **C.** The heatmap displays 308 ERV families (rows) enriched in either DLX5 or GATA3 ChIP-seq profiles. Heatmap showing the Spearman’s rank correlation matrix of the datasets from the left panel, based on adjusted p-values for enrichment within the 308 ERV families. The three clusters, separated by white lines, represent k-means clustering of the adjusted p-values. **D.** The heatmap displays the selected 16 ERV families (rows) enriched within GATA3 and DLX5 ChIP-seq profiles. These ERV families represent the top candidates, exhibiting either enhancer signatures (P300, MED1, H3K4Me1) or promoter signatures (H3K4Me3). Additionally, the MLT1 family is included as a negative observation, as it shows DLX5 enrichment but lacks the corresponding histone marks required to be classified as an enhancer or promoter. **E.** Phylogenetic origin of selected ERV families shown on Fig. 1C (right panel). The figure shows the approximate time of ERV fixation. Note that all shown ERVs were fixed during primate evolution, with the exception of the LTR family Mammalian LTR transposon 1 (MLT1). In particular, the *MLT1F2* and *MLT1A1* families, which are members of the Mammalian apparent LTR retrotransposon (MaLRs) superfamily, were already fixed before the divergence of primates from other mammals [120].

From this dataset, we focused on ERV families enriched in accessible genomic regions with enhancer or promoter characteristics and in the binding profiles of both GATA3 and DLX5. We identified a total of 308 ERV families that showed significant enrichment in at least one of the analyzed ChIP-seq profiles, with most being positive for the H3K9Me3 mark and none for the H3K27Me3 mark (Fig. 1B-C). Among the 308 ERV families, 144 showed significant binding of either GATA3 or DLX5, with 97 enriched for GATA3 binding and 131 for DLX5, and 84 families shared between them, despite the GATA3 ChIP-seq being performed in TSCs and the DLX5 ChIP-seq in EVT cells. These shared ERV families were also enriched with P300, ATAC-seq, MED1, H3K4me1, and H3K27ac marks (82% of DLX5-GATA3 shared peaks), which are hallmarks of active enhancers. These ERVs include some that are evolutionarily older, such as the MLT1 families. Their transcriptional regulatory activity as placental enhancers and their dysregulation in pre-eclampsia have recently been reported [89]. We surmise that, while pre-eclampsia-dysregulated TFs targeting ERVs exhibit strong enhancer potential, only a few ERV families were enriched within the promoter repertoire of the trophoblast lineage. Our analysis corroborates previous observations showing higher promoter activity in the youngest RE families, such as L1_Hs, SVA, and LTR5_Hs (Fig. 1D).

Among all the TFs analysed, our findings specifically highlight TFAP2C, which exhibited exclusive co-occupancy with H3K4me3 deposition, suggesting it as a key factor in classifying ERVs with promoter activity. Interestingly, the LTR8A and LTR8B subfamilies showed different enhancer and promoter activity, with the enrichment of TFAP2C binding specifically within the LTR8B family indicating its promoter potential (Fig. 1D).

Together, the analysis suggests that GATA3 and DLX5, in conjunction with specific ERV-derived sequences, represent candidate loci for placental gene regulation during primate evolution [7] (Fig. 1D-E). Among the shortlisted ERV families, we identified LTR5_Hs, LTR8B, and LTR10D, which also met the criteria of being bound by the pioneer factor TFAP2. This approach ultimately led us to focus on LTR8B.

### The recently expanded Pregnancy Specific Glycoproteins (PSG) locus is rich in ERV-derived, potential regulatory sequences

Among the shortlisted ERV families, we observed an enrichment of LTR8B elements at the pregnancy-specific glycoprotein (PSG) gene array [30], which has previously been associated with the human-specific pregnancy disease pre-eclampsia (PE), albeit with rather contradictory data [45]. By contrast LTR5_Hs, the other top candidate, had no placenta-specific genes in close proximity. The LTR8B elements are associated with all members of the PSG gene array (Fig. 2A), consistent with them having been amplified along with the genes during the expansion of this locus. TF binding, histone marks, and chromatin accessibility identify LTR8B as a prominent cis-regulatory element for its corresponding PSGs (Fig. 2A). Among the co-amplified elements, we also find MER65 (Fig. 2A) located at the 3’ ends of PSGs. The enrichment of ERVs at the PSG locus as well as its potential association with PE made the PSG locus our choice for our study. Specifically, as the PSG genes have no orthologs in non-primates [90], and have no clear transcription regulatory sequences [91, 92], we hypothesized that LTR8B and MER65, found exclusively in anthropoid primates (Additional File 2: Fig. S1), might provide regulatory signals for the PSG genes, and the PSG genes themselves contributed to the evolution of primate placentation.

**Fig. 2.**
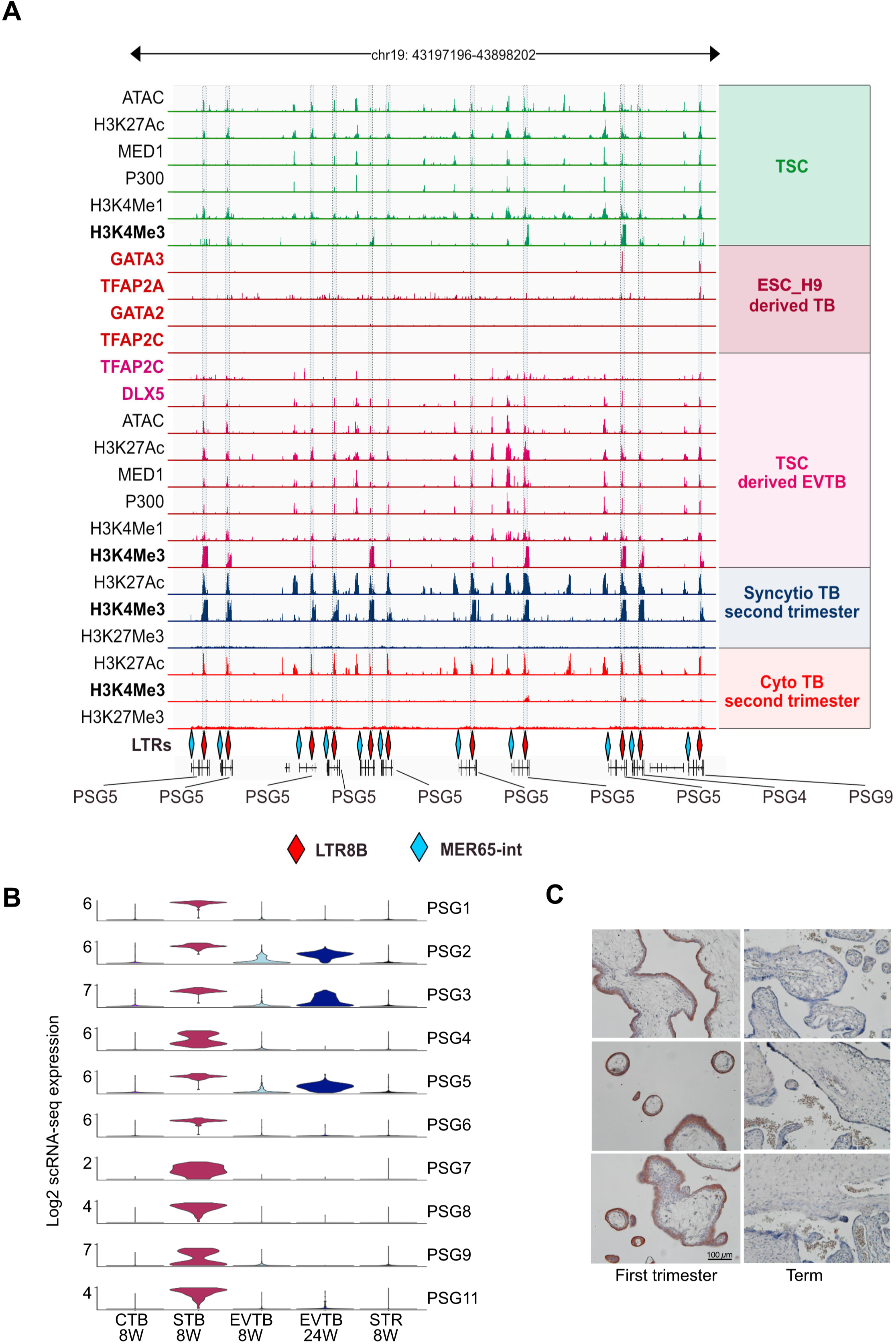
PSG family analyses. **A.** Overview of the chromatin landscape of the human PSG gene array, which encodes 10 protein-coding genes. An Integrative Genome Viewer (IGV) was used to visualize the normalized ChIP-seq signals of multiple histone modifications, transcription factors (TFs), enhancer markers (P300 and MED1), and ATAC-seq data across the entire human PSG gene cluster. The positions of two ERV elements, LTR8B and MER65, co-amplified with the PSG genes, are indicated. The data were sourced from: TSC and TSC-derived EVTB [101], ESC_H9 [88], Cytotrophoblast (CytTB) and Syncytiotrophoblast (SynTB) [121]. **B.** Differential cell type specific expression patterns of PSG genes in distinct trophoblast populations (cytotrophoblasts (CytTB), syncytiotrophoblasts (SynTB), and extravillous trophoblasts (EVTB)) of first and second trimester placenta samples; Single cell (sc)transcriptome, SMART-seq2 [93]. **C.** Representative immunohistochemistry staining of PSG9 in placental villi of first trimester (left panels) and term pregnancy (right panels); PSG9 Ab 8 μg/ml.

### LTR8B at the PSG locus defines cell type and development stage specific expression

While LTR8B is repeated at the PSG array, chromatin marks and the TF binding signals are not equally distributed along the LTR8B copies at the PSG array. Therefore, we performed some family-wide analyses. We considered transcription factors with known affinity for ERV derived sequences (GATA2/3 [7]) as well as placentally important factors (e.g. DLX5, TFAP2A and TFAP2C) ChIP-seq analysis via data mining (Fig. 2A). We found evidence that LTR8B elements might serve as binding platforms for DLX5 and TFAP2A. These factors could potentially be added to the list of LTR8B-binding transcription factors relevant to the placenta, which also includes FOSB and JUNB, however, our positive result for TFAP2C conflicts with previous analyses [30]. We confirm binding with GATA3 but add that there is no evidence for GATA2 (Fig. 2A). These TFs are expressed at different ratios in the three trophoblast lineages, suggesting that the unique combination of these factors potentially defines lineage specific expression regulation of the PSG members. Indeed, data mining of the placental single-cell transcriptome from the first trimester of pregnancy [93] reveals that while the different PSG family members have characteristic expression profiles, they are all predominantly expressed in the syncytiotrophoblast (STB) [93] compared to the extravillous trophoblast (EVTB) (Fig. 2B). Interestingly, PSG9 stands out in this analysis: the LTR8B element associated with PSG9 is distinct in being enriched with binding sites for a specific subset of transcription factors (TFs) among all those tested, including GATA3, DLX5, TFAP2A, and TFAP2C (Fig. 2A).

Therefore, to further investigate cell-type as well as stage-specific expression at the protein level, we performed immunohistochemistry using an antibody specific to PSG9 (Additional File 2: Fig. S2) on human placenta tissues (first and third trimester samples) (Fig. 2C). In agreement with the single cell transcriptome data (Fig. 2B), the staining detected PSG9 primarily in STB (Fig. 2C). Furthermore, term placenta showed weak staining for PSG9, when compared to first trimester (Fig. 2C).

### The family member PSG9 is dysregulated in PE

Abnormal expression levels of several family members (e.g. PSG1, PSG7, PSG11 and PSG9) have also been reported in PE, however, their contribution to this pathological pregnancy is rather controversial [45, 59]. To clarify, we analysed various PE datasets (Additional File 3). First, we performed transcriptome analysis (bulk RNA-seq, Additional File 4) on our set (10 PE; 8 healthy) of trophoblast samples (Charité, Berlin cohort). Transcriptome analysis revealed the upregulation of two family members (e.g. PSG11 and PSG9), whereas the transcriptional changes of the other PSGs were not significant (Fig. 3A). Increased expression of PSG9 and PSG11 was confirmed by RT-qPCR (Fig. 3B), levels of which correlated with clinical markers (Fig. 3C).

**Fig. 3.**
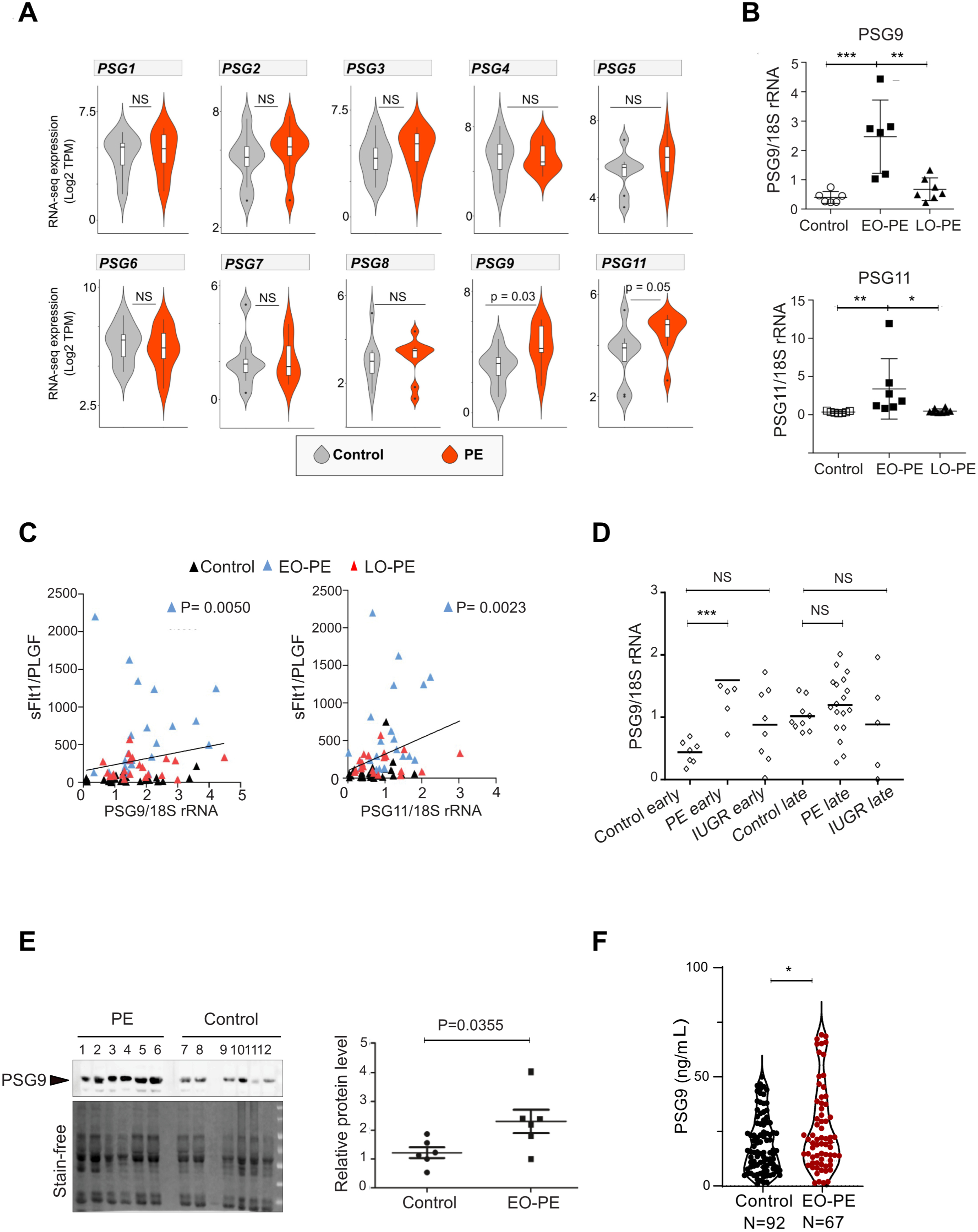
The PSG family member PSG9 is upregulated in the human pregnancy disorder, PE. **A.** Violin plots illustrating the expression dynamics of PSG genes in healthy pregnancy and in patients with early-onset pre-eclampsia (EO-PE). RNA-sequencing analysis (8 healthy controls vs 10 EO-PE) (Charité-Berlin cohort). **B.** qRT-PCR validation of the dysregulated expression of PSG9 and PSG11 is dysregulated in EO-PE. **C.** Correlation of dysregulated expression of PSG9 and PSG11 with clinical PE biomarkers. The placental expression of PSG9 and PSG11 was plotted against the ratios of the “anti-angiogenic” sFlt1 and “pro-angiogenic” PlGF (sFlt1/PlGF) concentrations in maternal serum. The expression values were normalized to 18S. **D.** Dysregulated expression of PSG9 is specific to EO-PE (Charité-Aachen cohort). qRT-PCR analysis of PSG9 mRNA expression in PE, Intrauterine Growth Restriction (IUGR) patients and healthy controls (early and late). The mRNA expression was normalized to 18S rRNA. Unpaired t test. Note that only the differences between the early onset PE group and the healthy control group were significant *** P= 0.0007. **E.** Upregulated PSG9 protein expression is detectable in the maternal serum of EO-PE patients (Manchester cohort, samples collected between 28-30 weeks). Left panel: Western blot (top) (ab64425, Abcam), stain free blot-loading control, quantification (bottom). Right panel: Quantification of the Western blot using Image Lab™ software (Bio-Rad). The bar shows the mean ±SEM (unpaired t-test). **F.** Dysregulated expression of PSG9 is detectable in the maternal serum. Sandwich ELISA for specifically detecting PSG9 (NBP1-57656, Bio-Techne, Germany). Using our sandwich ELISA, we analysed the levels of PSG9 in the serum of 67 patients affected by EO-PE and 92 uneventful pregnancies (Hungary cohort, term) [94]. The sandwich ELISA supported the hypothesis that the maternal circulating PSG9 levels were higher in EO-PE patients (P < 0.05) than that in control pregnancy. Mann Whitney U test.

It is important to note that gestational age, maternal blood pressure, BMI, and birth weight can all influence placental gene expression. These covariate effects have been extensively analysed in our previous work [38], where the gene sets associated with each clinical parameter were systematically reported.

Significant upregulation of PSG9 was specific to early-onset PE (EO-PE) *versus* late-onset PE (LO-PE) and an unrelated pregnancy disorder IUGR (intrauterine growth restriction) (Fig. 3D). Using an antibody that exclusively recognizes PSG9 (Additional File 2: Fig. S2), we asked whether upregulated PSG9 could be also detected at the protein level in the maternal serum. Analysis of PE and healthy samples from the Manchester cohort suggested an elevated level of PSG9 in PE serum samples (Fig. 3E). To enable precise monitoring maternal PSG9 circulating levels, we established a PSG9-specific sandwich ELISA, using a validated PSG9-specific antibody, and analysed the levels of PSG9 in the serum of 67 patients affected by EO-PE and 92 uneventful pregnancies (Hungary cohort, term) [94] (Fig. 3F). The sandwich ELISA supported the hypothesis that the maternal circulating PSG9 levels were higher in EO-PE patients (P < 0.05) than that in control pregnancy (Fig. 3F).

### The LTR8B elicits placenta-specific cCREs property at the PSG9 locus that potentially interacts with multiple genomic loci

To find a possible explanation to the relative uniqueness of PSG9 among PSGs, we performed an in-depth analysis on the PSG9 locus. We began analysing ERV-derived sequences that had been co-amplified with the PSG genes (e.g. LTR8B and MER65) (Additional File 2: Fig. S3). The PSG9 locus appears to produce multiple alternative transcripts, as evidenced by alternative splicing and RNA-seq analyses (Fig. 4A). To validate the regulatory potential of LTR8B at the PSG9 locus, we employed a luciferase reporter assay on a trophoblast cell line that is commonly used to study syncytialization (BeWo) [95]. In the assay, in addition to LTR8B, we systematically tested multiple TE-derived sequences, amplified from the PSG9 locus (Fig. 4B). In the reporter assay (Additional File 2: Fig. S4), LTR8B exhibited the most robust trophoblast-specific luminescence signal compared to the other TE-derived elements and the negative control. This supports its regulatory potential as a cis-regulatory element (CRE) to drive gene expression.

**Fig. 4.**
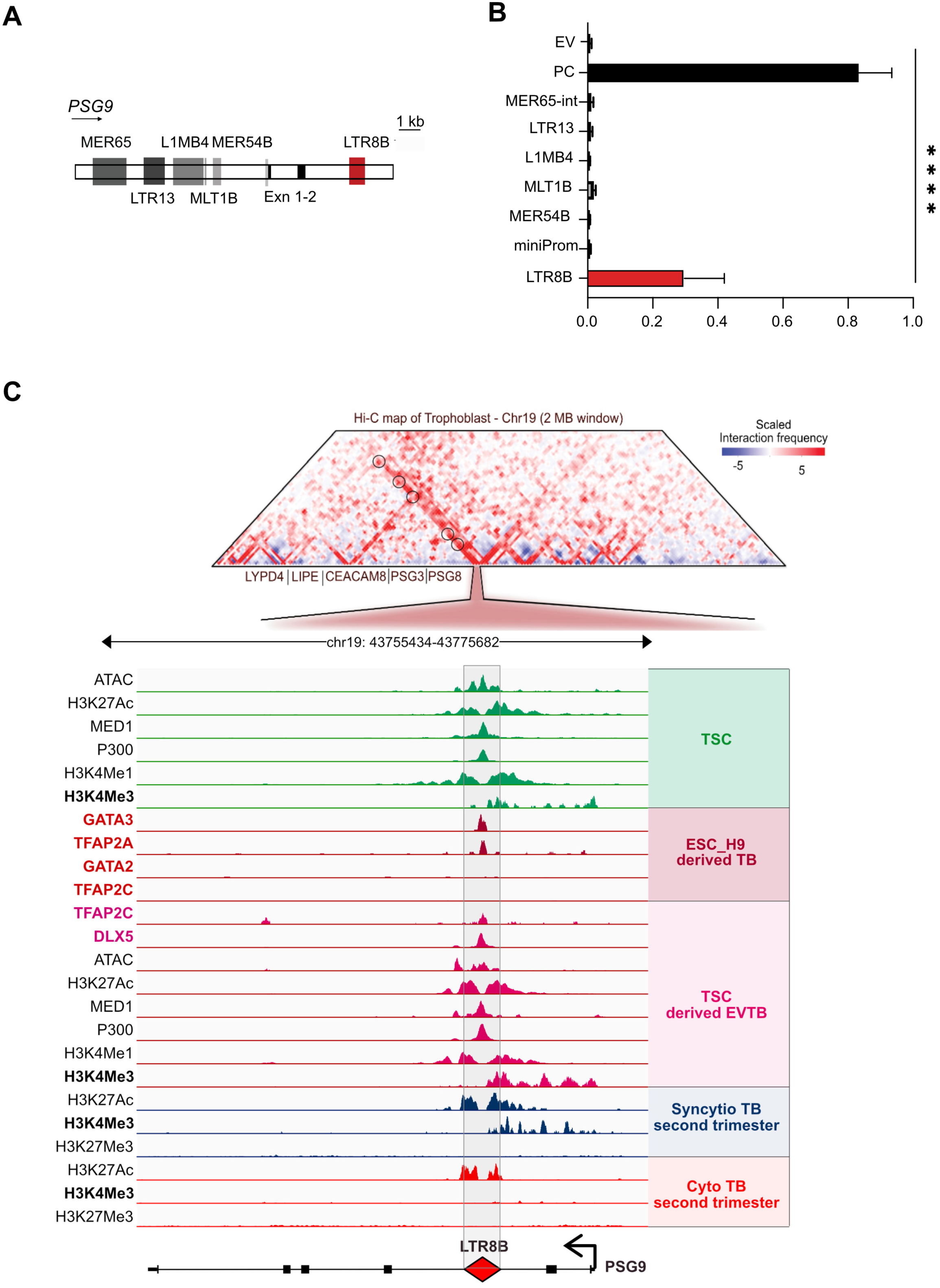
LTR8B is a trophoblast-specific cCRE at the PSG9 locus. **A. The PSG9 locus produces multiple alternative transcripts.** Genome browser snapshot showing characteristics of the PSG9 locus: GENCODE Genes track (version 48); Repeated elements by RepeatMasker; Alternative splicing graph from Swiss Institute of Bioinformatics. The underneath RNA-seq tracks represent the expression at individual exon resolution from various trophoblast lineages (data source: GSE204722). **B.** (Upper) Schematic figure showing the position of LTR8B and the upstream elements at the PSG9 locus tested in the reported assay shown on Fig. 4B). Exn, exon. **C.** Reporter assay to validate LTR8B/PSG9 as a functional enhancer. The enhancer activity was evaluated based on the luciferase reporter assay. EV; empty vector, PC, positive control, SV40 enhancer. N = 6 technical replicates, mean ± SD. One-way ANOVA followed by multiple comparisons. Empty vector vs. LTR8, **** *P* < 0.0001. **D.** Chromatin interaction profiling of the *PSG9* gene region. (Upper) Hi-C data visualization showing the chromatin interaction of the LTR8B element in hESC_H1-derived trophectoderm cells (GRCh38/hg38). Data mining of [98, 122], visualized by the tool ‘3D-Genome Interaction Viewer & database (3DIV)’ (http://www.3div.kr/hic and http://www.3div.kr/capture_hic) [74]. (Lower) UCSC genome browser visualisation of the chromatin features of the LTR8B at the PSG9 locus with the same set of data shown in Fig. 2A. TSC, trophoblast stem cell.

While, LTR8B sequence was previously shown to function as an enhancer [7, 30, 96], the promoters of PSG genes remain poorly defined [91, 92]. In-depth analysis revealed H3K4Me3 deposition and TFAP2C binding, key determinants of promoter activity, at the LTR8B locus (Fig. 1D). To further clarify whether transcription initiates within the LTR8B element, we also analysed CAGE and raw RNA-seq aligned reads (Additional file 2: Figure S5) as well as ATAC-seq data (Fig. 4D). Collectively, these analyses suggest that, although LTR8B displays chromatin features typical of active promoters, PSG9 transcription is predominantly initiated from its canonical promoter.

We next integrated the multi-omics data with Hi-C and promoter capture profiling (PC Hi-C) data from trophoblast cells [97, 98]. This analysis revealed another noteworthy feature of the PSG9 locus: the LTR8B/PSG9 region engages in interactions with several nearby genomic loci, likely mediated by active chromatin loops (Fig. 4D and Additional File 2: Fig. S6-S7). Among the potential interacting loci, we identified other PSG family members (PSG3 and PSG8), as well as neighbouring genes LYPD4, LIPE, and CEACAM8 (Fig. 4D), suggesting the LTR8B-derived cCRE exerts regulatory effects beyond the PSG array.

### LTR8B/PSG9 is required for syncytiotrophoblast identity and governs the process of syncytiotrophoblast differentiation

To decipher if cCRE LTR8B is sole and the CRE at the PSG9 locus, we carried out a CRISPR/Cas9 mediated deletion of LTR8B. To knock out (KO) LTR8B, we used two different pairs of guide-RNAs flanking the element and monitored the expression of PSG9 in multiple homozygous clones (Fig. 5A and Additional File 2: Fig. S8A). We could not generate a single viable TSC colony following LTR8-KO. Notably, PSG9-LTR8B exhibits chromatin features associated with active transcription in hTSCs, suggesting that this locus may exert an additional regulatory role at the stem cell stage, potentially affecting trophoblast self-renewal or survival which could explain the absence of viable KO-hTSC clones. Following several unsuccessful attempts to screen KO-LTR8B human trophoblast stem cells (hTSCs), we conducted the KO assay in BeWo cells. The five independent KO-clones exhibited downregulated PSG9 transcription levels (Fig. 5A), validating that LTR8B is the key CRE of PSG9 expression. Clone LTR8B KO P6-B4, which had almost undetectable PSG9 expression, was selected for downstream experiments (Fig. 5A).

**Fig. 5.**
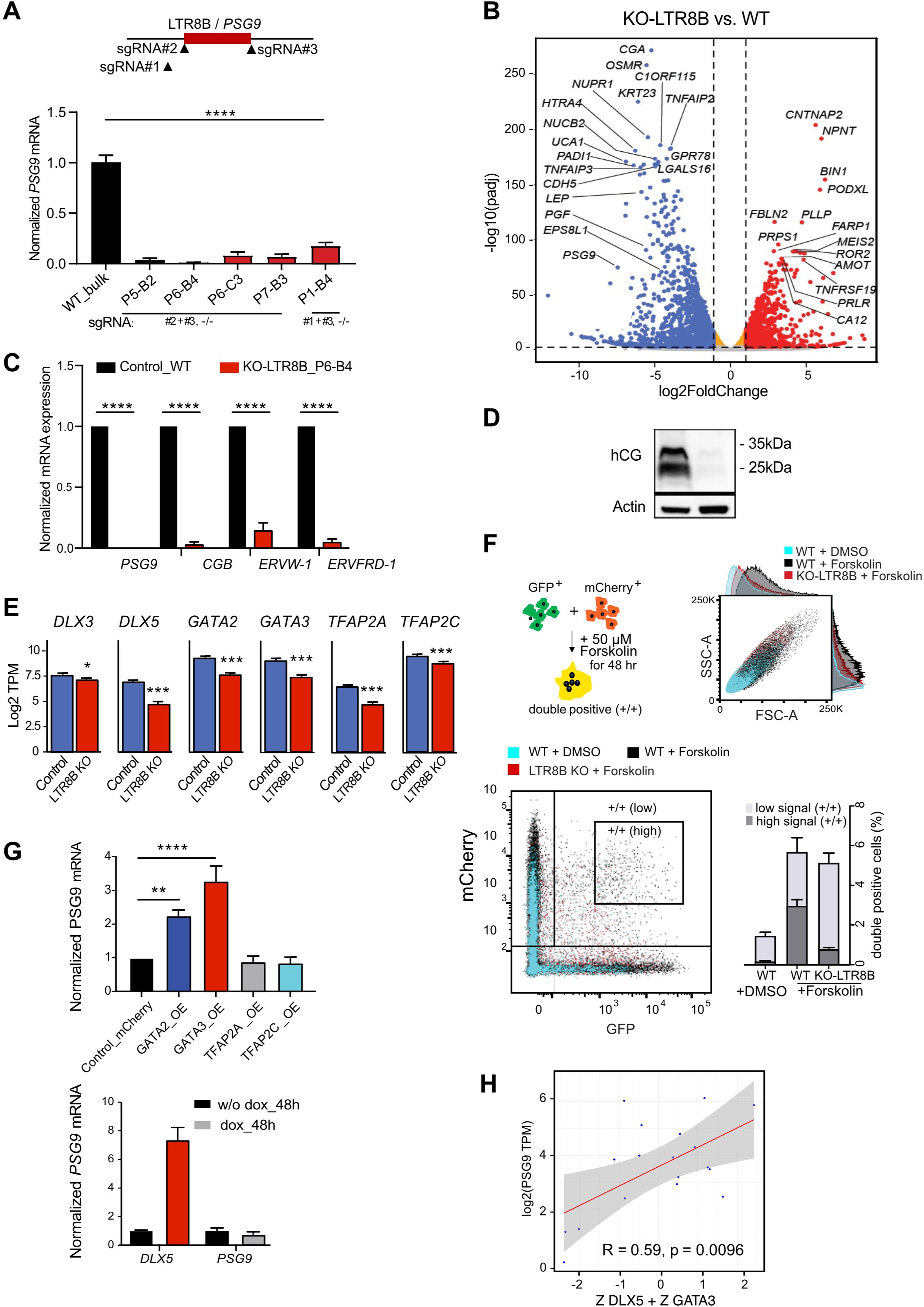
The LTR8B is the CRE at the PSG9 locus is essential for syncytiotrophoblast differentiation and identity. **A.** CRISPR/Cas9-directed deletion of the LTR8B putative enhancer at the PSG9 locus. Experimental design showing the knockout (KO) strategy. Two different pairs of sgRNAs targeting the LTR8B were introduced and mRNA expression of PSG9 in the homozygous KO colonies was examined (normalized (against wild type bulk cell population). The designations P5-B2, P6-B4, P6-C3, P7-B3, and P1-B4 refer to individual CRISPR-Cas9-mediated LTR8B KO clonal cell lines. Each label represents a different clone, isolated and expanded for downstream validation and analysis. N = 3, mean ± SD. One-way ANOVA followed by multiple comparisons **** *P* < 0.0001. **B.** Volcano plot visualizing the DEGs in the LTR8 KO BeWo cells (clone P6-B4) (N=3 per group) compared to the wide type control. The X axis represents the differential expression levels transformed into the log_2_ fold change and the y axis represents the adjusted p-values attained by the Wald test with correction for multiple testing using the Benjamini and Hochberg method. The log2 fold change cutoff = 2; the adjusted p-value cutoff = 0.000001. **C.** Expression of syncytialization signature genes in KO LTR8B BeWo cells (RT-qPCR, 48 h post-Forskolin treatment). N = 6 and 8 independent replicates for the KO LTR8 colony (P6-B4). Mean ± SD. Unpaired t test. **** *P* < 0.0001. **D.** Western blot showing the production of hCG protein in the *in vitro* BeWo syncytialization assay in KO-LTR8B/PSG9 cells (48 hr post-forskolin treatment). The left lane shows the expression of hCG in wild-type (WT) cells, while the right lane shows the expression in cells derived from the LTR8B KO P6-B4 clone. **E.** Negatively regulated trophoblast specific transcription factors (TFs) in KO-LTR8B/PSG9 BeWo cells (48 hr post-forskolin treatment, RNA-seq). **F.** Forskolin induced cell fusion assay by co-culturing the GFP+ and mCherry+ expressing control (LTR8 KO_P5-B2) and KO LTR8B (LTR8 KO_P6-B4) BeWo cells. Strategy to monitor forskolin-induced cell fusion of BeWo cells (syncytialization) by FACS. The GFP+ and mCherry+-expressing cells are co-cultured and then treated with forskolin (48 h) (Upper left). The forskolin treatment induces the cell fusion process. Upon forskolin treatment, the size of the cells increases (SSC-A/FSCA), and the larger syncytium (Upper right) can be identified as double-positive, GFP+/mCherry+ fluorescence by FACS analysis (Lower left). FSC-A, forward scatter area; SSC-A, side scatter area. Each dot represents a single cell. The cells with high GFP+/mCherry+ signals (syncytium) are highlighted in a rectangular box. Quantification of double positive cells. (Lower right). Note the relative decreased number of the double positive (high signal +/+ cells). N = 3 independent replicates, mean ± SD. Paired t test. WT high signal vs. KO-LTR8 high signal, ** *P =* 0.004. **G.** (Upper panel) PSG9 transcription responds to the ectopic overexpression (OE) of trophoblast specific TFs, GATA2 and GATA3 (qRT-PCR) in BeWo cells. N=3 technical replicates, expression of ectopic TFs was validated by Western blot. N = 3 independent replicates, mean ± SD. Ordinary one-way ANOVA followed by multiple comparisions. Control vs. GATA2 OE, ** *P* = 0.0011; control vs. GATA3 OE, **** *P* < 0.0001; control vs. TFAP2A/C not significant. (Lower panel) RT-qPCR analysis of PSG9 expression upon DLX5 induction with Dox treatment (48 h) in a Doxycycline inducible DLX5 expressing BeWo line. **H.** GATA3 and DLX5 combined predict PSG9 expression. log2(RNA TPM) for (i) GATA3 and (ii) DLX5 levels are positively correlated with PSG9 levels, the later significantly so. The combined effect is, however, more significant. As GATA3 and DLX5 occur at different levels, we convert, for each, log2(RNA TPM) to Z scores: (observed – sample mean)/sample sd. We add the Z scores of the two genes for each sample (iii). Pearson correlation data given in each panel.

To investigate the possible mechanism behind the KO-LTR8B phenotype at the PSG9 locus, we performed a transcriptome analysis on KO-LTR8B syncytiotrophoblasts. This analysis revealed 1,257 differentially expressed genes (DEGs) (adjusted P value < 0.01). Among the most significant DEGs were chorionic gonadotropin alpha (CGA) and several beta subunits (CGB), which together form human chorionic gonadotropin (hCG), a key marker of syncytiotrophoblast (STB) lineage identity (Fig. 5C). Notably, the protein levels of hCG were nearly undetectable in KO-LTR8B cells (Fig. 5D). Moreover, the expression of a broad set of STB signature genes, including ERVW-1/Syncytin 1, ERVFRD-1/Syncytin 2, LGALS16 (galectin 16), and GDF15 (Growth Differentiation Factor 15), was significantly impacted (Fig. 5B and Additional File 2: Fig. S8B). RT-qPCR validation confirmed a reduction of ERVW-1/Syncytin 1 and ERVFRD-1/Syncytin 2, which are essential for syncytiotrophoblast fusion and are known to be critical to STB development (Fig. 5C). These findings indicate that knocking out LTR8B at the PSG9 locus leads to the loss of STB lineage identity.

Additionally, several trophoblast-specific transcription factors, such as DLX5, GATA2, GATA3, TFAP2A, and TFAP2C (Fig. 5E), were downregulated, potentially contributing to the loss of STB characteristics in KO-LTR8B/PSG9 cells. The knockout of LTR8B/PSG9 also disrupted the canonical forskolin-stimulated cAMP signaling pathway, evidenced by the strong downregulation of RAPGEF4 (EPAC2), a key regulator of processes like cell adhesion and migration [99]. Furthermore, transcription factors critical for syncytiotrophoblast formation, such as STAT5B [100] and GCM1 (Additional File 2: Fig. S8C), were also downregulated. In contrast, transcription factors associated with trophoblast lineage specification and cell fate determination, including CDX2 and TEAD4, were upregulated (Additional File 2: Fig. S8D). These factors likely promote self-renewal over differentiation, acting antagonistically to those required for syncytiotrophoblast differentiation during early trophoblast development. Collectively, these findings suggest that the knockout of LTR8B at the PSG9 locus interferes with the forskolin-induced differentiation process by disrupting its role as a cis-regulatory element (CRE), ultimately resulting in the loss of lineage identity in KO-LTR8B/PSG9 cells.

Knocking out LTR8B also affected the expression of neighbouring PSG loci (e.g. PSG2, 3 and 4). (Additional File 2: Fig. S8B). However, RNA-seq read alignment across the PSG locus revealed that, although the expression of these PSGs was reduced, reads still mapped across their exons (Additional File 2: Fig. S9). This suggests that the LTR8B element at the PSG9 locus primarily acts as a local enhancer for neighbouring PSGs rather than off-target editing effects being responsible for the reduction.

To demonstrate the significance of the deletion of LTR8B/PSG9 on the syncytialization (multinucleation) process, we performed a forskolin-induced cell fusion assay by co-culturing the GFP+ and mCherry+ expressing wild type control and KO LTR8B (KO-LTR8B/PSG9_P6-B4) BeWo cells. Following the forskolin treatment, we monitored both the cell size and the percentage of double-positive (GFP+/mCherry+) cells by fluorescence-activated cell sorting (FACS) (Fig. 5F). In line with the cell fusion process, the size of the control (WT) cells became larger (SSC-A/FSCA), and the number of fused cells (double-positive, GFP+/mCherry+) were higher compared to the WT vehicle-treated cells upon forskolin treatment. Compared to the forskolin-treated WT cells, the LTR8B/PSG9 KO cell population displayed a smaller cell size, and a significantly reduced number of highly fluorescent double-positive cells (∼5-fold lower) (Fig. 5F), suggesting that the KO-LTR8B/PSG9 cells were compromised in their ability to form a syncytium.

Finally, to determine whether PSG9 expression could compensate for the absence of LTR8B, we conducted rescue experiments involving the overexpression of both membrane-bound and secreted PSG9 isoforms (PSG9-201, PSG9-202 and both together) in an LTR8B knockout background. Notably, none of the isoforms, either individually or in combination, restored the expression of STB marker genes (see Additional File 2: Fig. S10). In line with this, our chromatin interaction analysis suggests that LTR8B may regulate additional genes within the PSG cluster as well as adjacent loci. Therefore, the phenotype observed upon LTR8B knockout is likely the result of coordinated regulation of multiple targets, rather than PSG9 alone.

### GATA3 and DLX5 levels correlates with the expression of PSG9

To further decode the regulation of the PSG9 locus, we focused on the set of trophoblast-specific transcription factors (e.g. GATA2/3, TFAP2A/C, TFAP2C and DLX5) that bind the LTR8B/PSG9 platform (Fig. 2A), and were also down-regulated in KO LTR8B (Fig. 5E). To dissect their potential regulatory role, we co-expressed individual TFs with the LTR8B/PSG9 luciferase reporter construct in BeWo cells, and assayed for luciferase expression. The assay revealed that the ectopic expression of both GATA3 and TFAP2A TFs can drive the LTR8B/PSG9 CRE to express the reporter (Additional File 2: Fig. S11A). To add a chromatin context to the analysis, we used RT-qPCR to monitor the transcription of PSG9 in BeWo cells, ectopically co-transfected with the expression constructs of various TFs. In addition, we determined PSG9 level in a stable line, where we could induce DLX5 expression. While the overexpression of TFAP2A or DLX5 had no effect, the expression of PSG9 was increased by the overexpression of GATA2/3 (Fig. 5G). In a similar assay, we tested PSG1, a randomly selected other member of the PSG family; however, PSG1 expression was not affected by the overdose of any of the tested TFs indicating some degree of specificity of regulation of PSG9 by these moieties (Additional File 2: Fig. S11B-C). Taken together, GATA2/3 drives PSG9 expression via the LTR8B/PSG9 element.

Given that both GATA3 and DLX5 are upregulated in many instances of OE-PE [38], and that PSG9 expression specifically responded to overdosed GATA2/3 TF levels (Fig. 5G), we next asked whether their elevated expression correlates with PSG9 levels in patient data. In our EO-PE transcriptome dataset (N=10), we found that the combined expression of GATA3 and DLX5 is predictive of PSG9 expression (Pearson R=0.59; p=0.0096) (Fig. 5H), both showed positively correlations, although only DLX5 reached statistical significance (Pearson correlation, p=0.0405). Regarding GATA2/3, although the LTR8B element at the PSG9 locus harbours predicted GATA motifs, our data do not indicate significant GATA2 occupancy at this site (Fig. 2A). To clarify this, we determined the regulatory networks governed by GATA2 and GATA3, which revealed only limited overlap in the trophoblast cells (Additional file 2: Figure S12A). Notably, DLX5 was included in the GATA3 regulome, whereas other DLX family members (e.g. DLX3 and DLX4) are associated with GATA2. Consistent with this, PSG9 levels decrease alongside DLX5 in DLX5-depleted EVTBs derived from TSCs [101] (see Additional file 2: Figure S12B). Taken together, these observations suggest several possible models. First, GATA3 may act directly on PSG9, whereas DLX5 exerts a context-dependent role in PSG9 control. In contrast, GATA2 may influence PSG9 indirectly, either by binding to distal enhancers within the PSG locus that physically interact with the LTR8B/PSG9 CRE hub in 3D, or by regulating intermediary TFs, such as the AP-1 components, which are directly recruited to LTR8B [30]. In this model, GATA2 overexpression would enhance PSG9 expression by activating a broader trophoblast-specific regulatory program, even in the absence of direct LTR8B binding.

### PSG9 has both secreted and membrane anchored protein isoforms defined by the presence or absence of MER65

Next, we investigated the possible significance of the MER65 elements, which are located at the 3’ ends of the PSGs and, like LTR8B, are also co-amplified with the PSG gene array (Fig. 2A and Additional File 2: Fig. S3). Embedded in MER65 elements (ERV1, HEPSI), we identified the annotated, core polyadenylation signal (AAUAAA) [102] at the 3’-end of PSGs (Additional File 2: Fig. S13A/B), suggesting that MER65 elements might provide a polyadenylation signal to PSGs, as proposed for other ERVs [26, 33]. To discern the functional consequences of the MER65-derived polyA signal, we focused on the PSG9 locus, which is characterized by annotated transcripts, both with and without inclusion of the MER65 element (Fig. 6A).

**Fig. 6.**
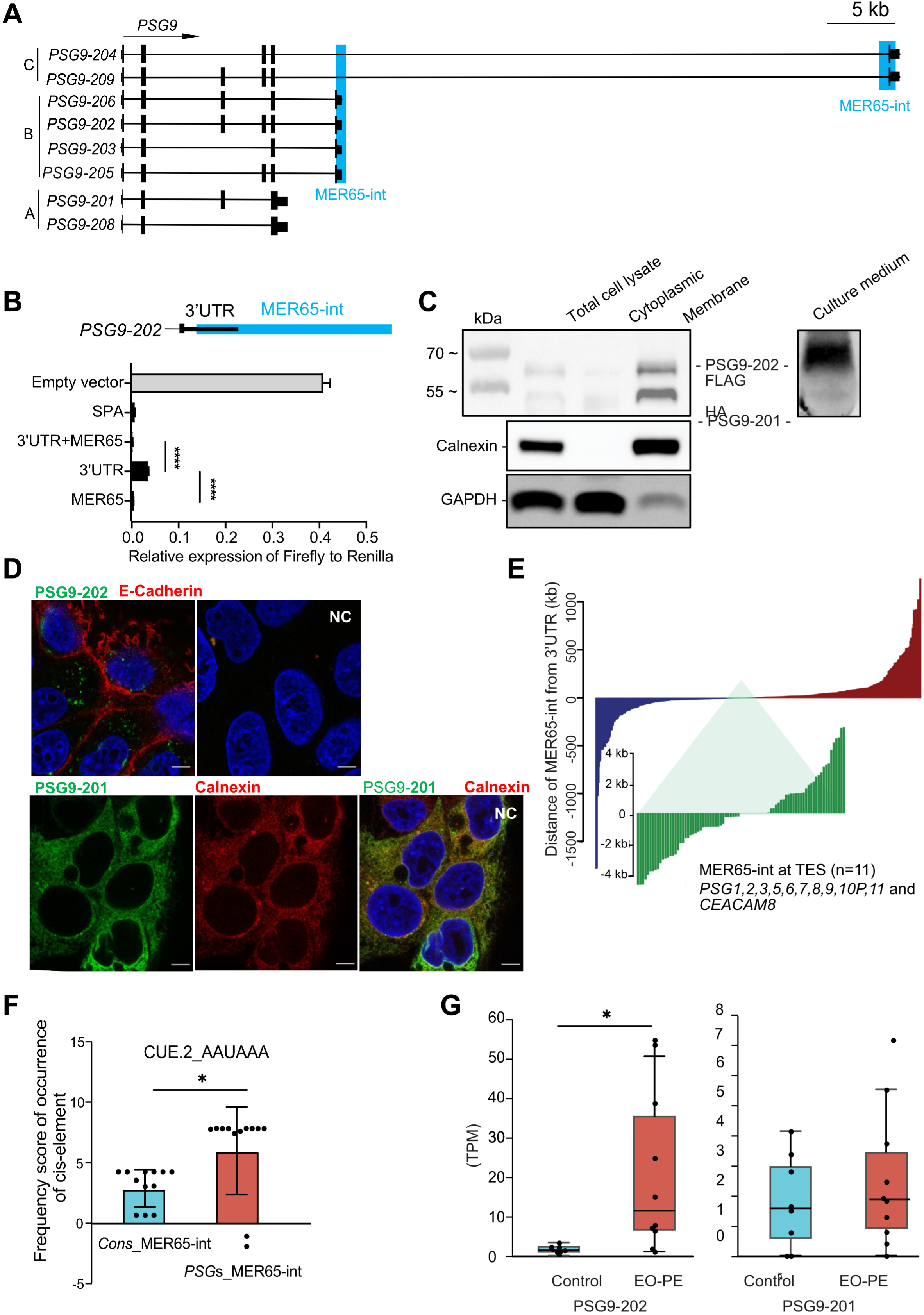
MER65 provides functional diversification of PSG9 transcripts via alternative polyadenylation. **A.** Structure of the various PSG9 isoforms. The annotated PSG9 isoforms can be classified in three groups, based on MER65-int (int - internal part, MER65 thereafter) (A,B,C). Group A has MER65, whereas groups B and C overlap with a proximal and distal MER65 element, respectively. For additional information on the isoforms see also Additional File 5. **B.** Validating the polyadenylation activity of the MER65 element using a reporter luciferase reporter assay. The polyadenylation strength of the MER65 element was determined as the relative expression ratio of Firefly versus Renilla luciferases. For the assay see also Additional File 2: Fig. S4) The MER65 element, the 3’ UTR upstream of the MER65 element and the combined 3’UTR + MER65 were tested. The empty vector served as a negative control, whereas a known synthetic polyadenylation signal (SPA) was used as a positive control. N = 6 technical replicates, mean ± SD. One-way ANOVA followed by multiple comparisons. **** *P* < 0.0001. **C.** MER65-directed alternative polyadenylation (APA) results in altered subcellular localization. Subcellar localization of the differentially tagged PSG9-201(-HA) (UniProt, Q00887-2) and PS9-202(-FLAG) (UniProt, Q00887-1) isoforms in a cell fractionation assay. The cell fractionation analysis visualized by Western blotting shows that the PSG9-202(-FLAG) was detectable in culture media (secreted), while the shorter isoform PSG9-201(-HA) was retained in the membrane compartment. Calnexin and GAPDH were used as fractionation markers. **D.** Differential subcellular localization of the PSG9-201(-HA) and PSG9-202(-FLAG) isoforms in trophoblasts. (Upper panels) Representative immunofluorescent co-staining of the PSG9-202(-FLAG) isoform and (E)-Cadherin (plasma membrane marker, marks the cell boundary). Note that PSG9-202(-FLAG) is detected in the intracellular space. (Lower panels) Representative immunofluorescent co-staining of the PSG9-201(-HA) isoform with Calnexin (ER marker). Note the overlapping staining signals between PSG9-201(-HA) isoform and Calnexin. NC, the cells stained with the secondary antibody as a negative control (marks the cytoplasmic compartment). Scale bar, 20 μm. **E.** Genome-wide analysis of the distance between MER65 elements and their host genes reveals that MER65 elements predominantly overlap with the 3’ UTR of PSG genes and CEACAM8 in humans. The Y-axis represents the distance of genes from MER65 elements within a range of 2.5 KB. The X-axis displays the genes, sorted in ascending order based on their distance to MER65 elements. **F.** Co-occurrence of the core upstream *cis*-element (CUE.2_AAUAAA) polyadenylation signal with MER65 sequences of the PSG genes. Paired t test, * *P* = 0.022. Each dot represents one MER65 sequence in *PSG* locus or the corresponding alignment in the consensus (cons) sequence. **G.** The PSG9-202 transcript is dysregulated in EO-PE samples when compared to healthy samples (10 EO-PE and 8 Healthy Control cells). P value: 0.016, FC:13,61, median: 1.35 (control) 11,12 (EO-PE); IQR 1,24 (control), 1,88 (EO-PE), whereas the PSG9-201 transcript was not differentially expressed 1.58 (control) 11,12 (EO-PE); IQR 2,38 (control), 2,51 (EO-PE).

To validate the potential of the PSG9-derived MER65 elements in generating transcription termination, we used a bicistronic reporter system [74], encoding for two different luciferase genes (Additional File 2: Fig. S14A). Although there are two MER65-derived elements at the PSG9 locus (Fig. 6A), we only tested the proximal copy of MER65, since the distal copy was not predicted to harbour a conserved polyA signal (Additional File 2: Fig. S13B). In the assay, we used MER65 alone, MER65 with an upstream 3’UTR sequence (3’UTR/MER65+) and the upstream 3’UTR without the MER65 (3’UTR/MER65-) (Fig. 6B). Further controls were synthetic polyA signal (SPA, positive) and the empty vector (negative). While the 3’UTR sequence (3’UTR/MER65-) generated some level of transcriptional pausing, the presence of the MER65 (3’UTR/MER65+) elevated pausing to the level of the positive control (SPA). Importantly, the PSG9-derived MER65 alone was capable of providing a nearly full transcriptional pausing effect (Fig. 6B), supporting that it functions as a polyA signal.

To assess whether the presence of MER65 contributes to alternative polyadenylation (APA), which may in turn define alternative protein function, we analysed the PSG9-201 and PSG9-202 isoforms using established methods [103]. These analyses predicted that PSG9-encoded protein variants differ in hydrophobicity (Additional File 2: Fig. S14B). For example, it is predicted that PSG9-201 will harbour a hydrophobic C-terminus and a transmembrane domain, whereas this region is absent in PSG9-202 due to the MER65-provided APA (Additional File 2: Fig. S14B-C, S15A and Additional File 6). To determine whether the presence or absence of the terminal exon influences cellular localization, we designed experiments to track two protein-coding PSG9 isoforms (e.g. PSG9-202 and PSG9-201), which either incorporate (+) or lack (-) the polyA-embedded MER65 element, respectively (Fig. 6C). We marked PSG9-201(-) and PSG9-202(+) protein variants with HA and FLAG tags, respectively, and stably co-expressed them in BeWo cells employing the *Sleeping Beauty* transposon system [69] (Additional File 2: Fig. S15B).

First, we subjected the cells to both subcellular fractionation and immunostaining. The fractionation assay detected both the FLAG-PSG9-202(+) and HA-tagged PSG9-201(-) protein variants in the membrane fraction. However, only FLAG-tagged PSG9-202(+) variant was detected in the culture media, suggesting that PSG9-202(+) is a secreted protein (Fig. 6C). Furthermore, immunostaining identified the FLAG-PSG9-202(+) variant in the intercellular space (Fig. 6D), while the membrane-bound, HA-labelled PSG9-201(-) isoform was intracellular and accumulated at the endoplasmic reticulum (ER) membrane of the cells, in agreement with a previous report but incorrectly referred to as PSG11w [80] (Fig. 6D). These observations are consistent with the possibility that PSG9 possesses both membrane-anchored and secreted protein isoforms. The extension of the transcript by alternative poly(A) addition signals could lead to alternative splicing events, providing more distal splice acceptor sites and resulting in the production of the secreted PSG9 protein isoforms (Fig. 4A and 6A).

To determine the generality of the phenomenon of MER65-mediated APA, we performed a genome-wide analysis of 912 MER65 copies across the human genome. Our analysis revealed that these elements are in proximity (< 50 bp) of the 3’end of genes exclusively at the PSG locus (with an exception of CEACAM8) (Fig. 6E) and that the core polyA signal (AAUAAA) is specifically enriched in MER65 elements located at the PSG loci (Fig. 6F). This is consistent with a co-option event of single MER65 element for post-transcriptional regulation by providing canonical or alternative polyadenylation signals particular to the gene cluster of PSG genes.

Nevertheless, the presence of both membrane-bond and secreted isoforms at the PSG9 locus tempted us to conduct a further bioinformatic analysis inside the PSG family members. Our deep learning pipeline revealed that PSG9 is the only PSG family member encoding for both membrane-anchored and secreted isoforms (Additional File 6). In contrast, all the rest of the family members have only the MER65-mediated secreted versions. Finally, we asked which PSG9 isoform is dysregulated in EO-PE. Our transcriptome analyses revealed that it is specifically the secreted PSG9-202(+) isoform whose upregulation can be detected in the trophoblast of EO-PE patients (Fig. 6G).

### Function of the PSG9 in the syncytiotrophoblast

The above data demonstrate the significance of the ERVs derived transcriptional sequences (e.g. LTR8B and MER65) in regulating PSG9. Next, we concentrated on the potential function of PSG9 itself. Previous reports associated PSG9 primarily with typical [101]B functions (i.e. affecting endothelial tube formation [59, 62, 63] or the invasion of the trophoblast [104], whereas its role in STBs, where it is primarily expressed, is less understood. To better understand its possible functions, we monitored PSG9 levels (by RT-qPCR) during the forskolin-induced STB differentiation process. We used again the BeWo cell fusion assay, mimicking trophoblast cell fusion [95, 105] and PCR primers specific for both membrane-bound (subgroup A, including PSG-201) and secreted (subgroup B, including PSG9-202) isoforms. Following forskolin induction, as the cells formed the multinucleated syncytium (up to 72 hours), PSG9 levels of both isoforms were gradually elevated (secreted and membrane-bound isoforms, ∼40-fold and ∼10-fold, respectively) (Fig. 7A). Intriguingly, single-cell transcriptome analysis revealed that PSG9 expression levels in STBs are comparable to those of established markers, such as CGA::CGB (chorionic gonadotropin) and Syncytin 1 (ERVW-1) (Fig. 7B), suggesting that PSG9 plays a key role in defining STB identity.

**Fig. 7.**
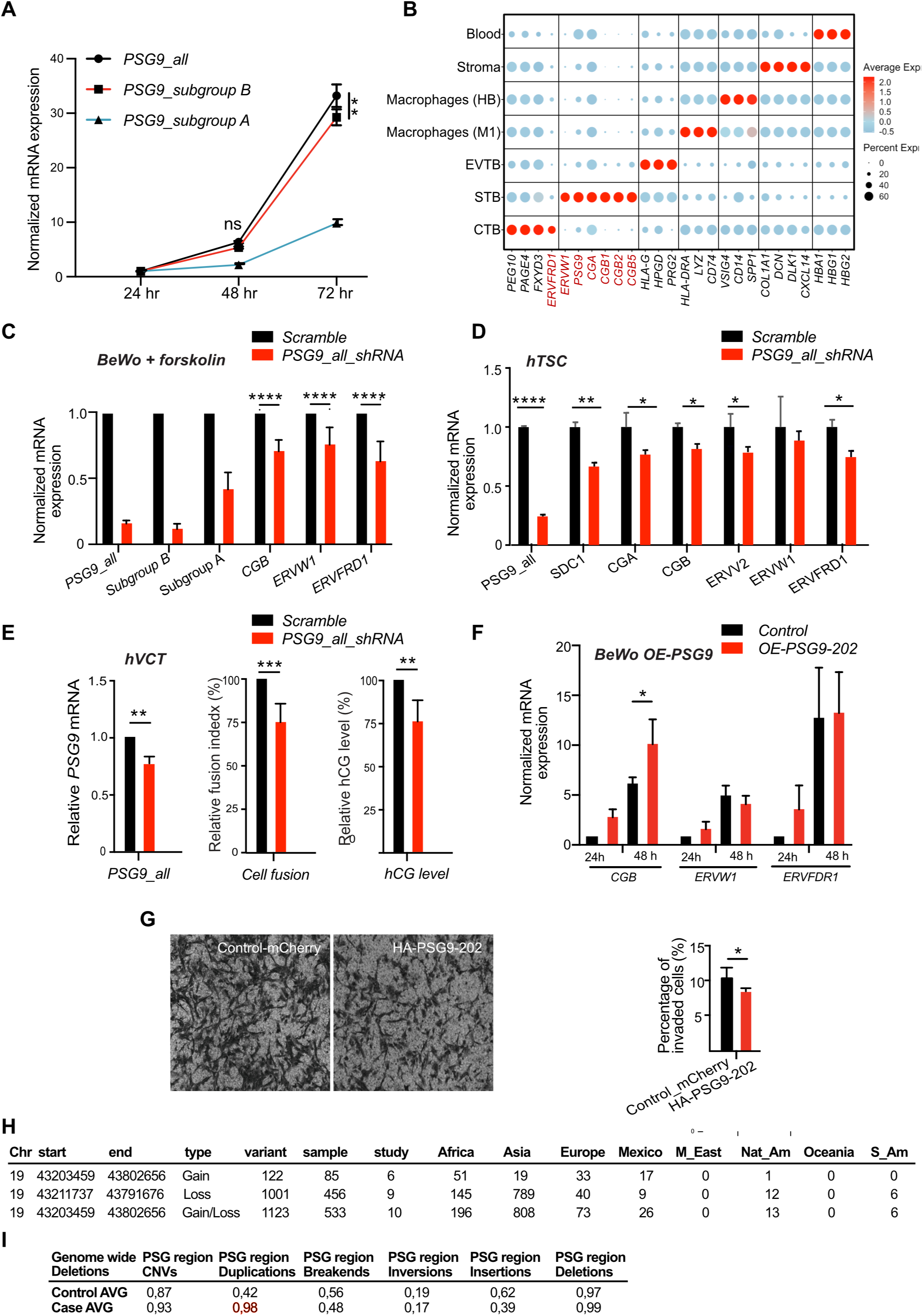
PSG9 function in STBs. **A.** RT-qPCR analysis of the PSG9 transcripts during the syncytialization process in BeWo cells (24, 48, 72 h); specific primers targeting all PSG9 isoforms, subgroup A (including the membrane bound PSG9-201) and B (including the secreted PSG9-202). N = 3 technical replicates, mean ± SD. Unpaired t test. *PSG9*_all vs *PSG9_subgroup B* at 72 hr, ** *P* = 0.002; *PSG9_subgroup B* vs *PSG9_subgroup A* at 72 hr, **** *P* < 0.0001. **B.** Dot plot illustrating the cell type-specific expression of the top marker genes for trophoblast subtypes (CTB, STB, and EVTB), macrophages (M1 and Hofbauer cells), stroma (decidual cells), and blood cells. Labels on the left of the plot indicate the cell types, while labels at the bottom denote the marker gene names. Gene names highlighted in red are of particular interest to this manuscript, including ERVW1 and ERVFRD1 (Syncytin 1-2), PSG9, and hCG genes (CGB1, CGB2, and CGB5). The colours represent the average log2 expression level, scaled to the number of unique molecular identifier (UMI) values per single cell. The colour scale ranges from light blue (lower expression) to red (higher expression), and the dot size is proportional to the percentage of cells expressing the corresponding gene. **C.** RT-qPCR analysis of expression of the syncytialization signature genes in KD PSG9 cells using shRNA constructs against both PSG9 subgroups (A and B), differentiated from BeWo cells (induced by forskolin). **D.** The effect of PSG9 depletion in the human trophoblast-stem-like cells (hTSCs) differentiated from ESCs [71]. **E.** Depletion of PSG9 affects cell fusion and hormone secretion negatively in the syncytium differentiated from human villous cytotrophoblasts (hVCTs). N = 5 **F.** Ectopic expression of PSG9-202 (secreted) in BeWo cells leads to increased CGB levels, whereas the expression of the syncytialization marker genes shows no difference (qRT-qPCR analysis, 24 h and 48 h following forskolin treatment). N = 3, mean ± SD. Two-way ANOVA followed by multiple comparisons. * *P* = 0.029. **G.** (Left panel) Representative image of PSG9-202 overexpression in the *in vitro* trans-well invasion assay in EVTB-like cell line (SGHPL-4) Scale bar, 200 μm (Right panel). Quantification of the *in vitro* transwell invasion assay. N = 6 (with four technical counts from non-overlapping microscopic images of each), mean + SD. One-way ANOVA followed by multiple comparisons (\**P* = 0.044). **H.** Copy number (CNV) analysis at the PSG locus reveals ethnic specific differences (1000 genome project). **I.** Structural variation (SV) analysis of the PSG locus region. Whole genome (gDNA) sequencing analysis of (21 healthy control and 76 PE patients, Oslo cohort). The analysis revealed sequence duplication (region of 1000 bp or larger) at the PSG region (p < 0.05; AVG set to 1).

To decipher the function of PSG9 in the STBs, we depleted PSG9 expression in BeWo cells using an RNAi strategy, and simultaneously subjected the cells to forskolin treatment, followed by transcription analysis. This approach detected the decreased level of PSG9 transcripts in the differentiated BeWo STB-like cells. Using RT-qPCR, we observed a significant knockdown (KD) effect (log2 FC = −3.3; adjusted p value = 0.03), when compared to the control (KD-scrambled) (Fig. 7C). We also detected significant downregulation of STB markers (e.g., CGB, ERVW-1/Syncytin 1, ERVFRD-1/Syncytin 2 (Fig. 7C). To further examine this, in addition to the BeWo cells, we determined the effect of PSG9 depletion in the human trophoblast-stem-like cells (hTSCs) [71], as well as in isolated primary human villous cytotrophoblasts (hVCTs). In the PSG9 depleted hTSCs, RT-qPCR detected significant decrease of the levels of CGA/B and multiple STB markers (e.g., ERVV2, ERVW-1/Syncytin 1, ERVFRD-1/Syncytin 2 and SDC1/Syndecan-1) (Fig. 7D), suggesting that the affected genes upon PSG9 depletion are consistent with typical STB identity. In addition, knocking down PSG9 in isolated human primary cells (hVCTs) had a negative effect on both the number of fused (multinucleated) cells and hCG levels in cellular assays (Fig. 7E). Overall, PSG9 contributes to determining the identity of STB and is involved in hormonal metabolism.

### Overdose of PSG9 affects trophoblast invasiveness

To dissect whether (and how) the elevated PSG9 protein itself might contribute the pathogenesis of PE, we overexpressed the PE dysregulated PSG9-202 (secreted) isoform which was driven by the CAGGS promoter. We consider levels of mRNA of marker genes and performed gain-of-function studies in trophoblasts.

Following forskolin-induced differentiation of BeWo cells, the overexpression of PSG9-202 resulted in an elevated level of *CGB* mRNA at 48 h post-treatment (p<0.05; Fig. 7F). Expression of other STB markers ERVW-1/Syncytin 1 and ERVFRD-1/Syncytin 2) showed no more of a tendency to upregulation at 48h than seen in control cells (Fig. 7F). These results are, at best, only weakly supportive of a role of PSG9 in modulating levels of markers.

Given that the secreted PSG9-202 isoform might even act on the surface of the neighbouring cells in the placenta, we also investigated the effect of PSG9-202 overdose in extravillous trophoblast (EVTB)-like cells. To this end, we overexpressed PSG9-202 in SGHPL-4 (EVT-like cell line [106]) and subjected the cells to transcriptome analysis. In response to the elevated level of PSG9-202 (logFC 7.72, p value = 3.86E-07), the analysis revealed a limited number of DEGs (number of genes N=9, p value < 0.01). The dysregulated categories included *organisation of the extracellular matrix* and *interferon signalling* (p value of the enrichment < 0.05). Notably, among the DEGs, we observed COL10A1 (Additional File 7) that in addition to its general structural functions of a collagen, has also been attributed to cell–cell interaction, tumour invasion, metastasis, and vascularization [107].

Following up on a recent study showing that the administration of recombinant PSG9 decreases trophoblast invasion in a 3D motility model (Swan 71 EVTB-like line) [104], we also investigated the effect of PSG9-202 overdose on SGHPL-4 cells in a trans-well invasion assay. Using the invasion assay, overexpression of the secreted variant of PSG9 resulted in reduced trophoblast invasiveness (Fig. 7G), suggesting that the upregulation of PSG9-202 might lead to insufficient trophoblast invasion.

### Evidence for CNV at the PSG locus

Thus far we have considered the PSG domain as though it was one fixed entity. However, likely owing to its repetitive nature, the PSG locus (chr19 (q13.2-q13.31)), with 47 structural variants (SVs), is among the most variable regions of the human genome [108]. The high copy number variation (CNV) of this locus is thought to be associated with pregnancy disorders, including PE [90, 109]. Copy-number deletion of PSG9 in this region was suggested to confer a risk for PE [65], but this could not be confirmed by other studies [45].

To consider whether PSG9 loss is consistently reported in PE patients, which would be contradictory to our finding of raised PSG9 levels, we reconsider the CSV issue in this genomic domain: chr 19: 43203459-43802656 region (PSG array). In the first instance we employ data from the 1000 genome project [110, 111] so as to better understand the global variation at the locus. Our analysis identified both loss and gain types of CNVs in different human populations (Fig. 7H). We observed higher ratios of loss to gain of copy numbers in the native American (12:1 ratio), African (2.8:1) and Asian (41:1) populations (Fig. 7H). In contrast, the analysis indicated an increase in the number of copies in the Mexican population (1:1.9) with only a small loss excess in Europe (1:1.2) (Fig. 7H). These results underscore the hyper copy number variability at this locus.

The relevance for such variation for the etiology of PE is unclear. To analyse genetic material collected from PE patients, we conducted whole genome sequencing (WGS) (76 PE and 21 healthy samples; Oslo cohort, European population). Importantly, our data fail to replicate prior claims of sequence loss being associated with PE [65], but accords with the above global CNV study. In contrast to the reported copy-number deletion (Asian) [65], our analysis performed on PE trophoblast samples (Oslo, European) detected a genomic duplication (Mann-Whitney U test, P < 0.05) at the PSG genomic region (Fig. 7I). How these effects modulate levels of PSGs, including PSG9 is unknown but worthy of follow-on scrutiny.

## Discussion

Given their placental co-option/expression and lineage-specificity, dysregulated ERVs are strong a priori candidates for human-specific pregnancy disorders, such as pre-eclampsia (PE) [30]. Intersection of genes dysregulated in pre-eclampsia and associated with RE mediated TF regulation led us to the PSG array. We identify that REs have contributed two important features enabling evolutionary innovations.

First, a key event in the evolution of the secretory capability of PSGs from their ancestral membrane isoforms was the evolution of the transcript that encapsulates MER65. In conjunction with alternative splicing, this enables polyadenylation of the novel form.

This accords with a recent suggestion that the evolution of STB differentiation elicits a widespread phenomenon of 3’UTRs shortening via APA, leading to secreted isoforms as an alternative to membrane anchored ones [112]. A parsimonious model is that MER65 inserted post the duplication of a membrane bound CEACAM (making the ancestral PSG), prior to PSG expansion. MER65 then enabled the evolution of a secreted isoform by shortening of the protein including loss of the hydrophobic membrane bound end while enabling polyA stabilization. This structure was then retained with array expansion defining the PSG family. By this route a secreted protein family evolved from a membrane bound ancestor. Whether PSG9’s membrane located version reflects an independent gain or the ancestral condition, is not transparent. Why the secreted form might be advantageous is also not transparent but a role in blocking adhesion receptors on maternal lymphocytes – comparable to the binding of CEACAM as membrane bound cell adhesion molecules – has been suggested [113]. While however maternal immune rejection of the foetus is considered a component of PE, such a model would more obviously predict that under-expression of PSGs would be associated with PE which is not universally observed.

However, the impact of MER65-int elements may extend beyond the modification of polyadenylation. Notably, the distal MER65 elements, which do not provide APA signals, overlap with distinct PSG9 isoforms (e.g. PSG9-204 and PSG9-209). These isoforms include an alternative exon that is absent from the other PSG9 isoforms, giving rise to predicted protein products with distinct AlphaFold-predicted tertiary structures (https://www.uniprot.org), suggesting potential functional divergence. Nevertheless, their biological relevance remains unclear and warrants further investigation.

The second key innovation permitted by REs (LTR8B) was in providing a binding platform for TFs and thus acting as CREs. At the point of duplication, the sequences of ancestral gene and duplicate would be identical and so the initial advantage – if any – of the duplication may have been a simple increase in PSG dosage. If that is the origin, it appears not to be what maintains the array. Our data instead supports that notion that different LTR8Bs provide varied binding platforms for TFs [30], each member of the PSG family being controlled by a specific set of TFs that define distinct expression patterns of PSGs. Whether this diversity reflects a sub-division of ancestral roles (sub-functionalization) or gain of novel roles (neofunctionalization) is not resolved. Given ongoing diversification and rapid evolution, neofunctionalization is more likely.

### PSG9s exceptionalism in normal and pathological placentation

While the above observations apply to the entire PSG gene array, the PSG9 locus is exceptional in several respects. First, PSG9 is the only family member that produces both secreted and membrane-bound isoforms (depending on the presence or absence of MER65 in the PSG9 primary transcript). Second, the LTR8B/PSG9 locus serves as a recruitment platform for a distinct set of TFs (e.g., GATA3, TFAP2A/C, TFAP2C, and DLX5). Third, LTR8B at the PSG9 locus not only regulates PSG9 expression but also anchors a greater number of chromatin loops than any other family member. LTR8B thus functions as both a local enhancer for nearby genes within these loops and as a cis-regulatory element driving PSG9 expression.

Although LTR8B exhibits promoter-like chromatin features and supports antisense transcription, the primary driver of PSG9 mRNA is likely its canonical promoter. However, RNA-seq read alignment following CRISPR–Cas9-mediated deletion of LTR8B at the PSG9 locus revealed a complete loss of PSG9 transcription, while other PSG family members retained detectable, though reduced, transcript levels. Importantly, the LTR8B/PSG9 locus also governs the canonical signalling pathway for trophoblast syncytialization in humans. Finally, PSG9 (secretory isoform PSG9-202) is the only member of the PSG family whose overdose is detectable in the blood of patients with early-onset pre-eclampsia (EO-PE).

The latter is potentially important as it suggests a (much needed) possible biomarker for EO-PE. While most PE cohorts come from symptomatic patients in the second or third trimester, there are signs that PSG9 levels could be elevated earlier in pregnancy [67]. Future research should systematically examine the predictive value of PSG9 in plasma samples collected before 12 weeks of gestation, which is the current cutoff for starting low-dose aspirin prophylaxis. Developing a reliable ELISA for PSG family members has proven challenging due to their structural similarities. Commercially available antibodies often lack validation and cross-react with other family members and certain CEACAM proteins [45, 68]. Despite these challenges, we are confident that our specific sandwich ELISA assay can accurately measure PSG9 levels in serum. This assay was rigorously tested using a mutant form of PSG9 (N-domain mutant), which further confirmed its specificity and robustness in detecting PSG9 without significant cross-reactivity.

Whether and how PSG9 overdose plays a causative role in PE remains unresolved. Further studies are also required to understand the role of the multiple isoforms, including those generated by the distant copy of MER65-mediated APA. A possible clue is provided by the fact of an imbalance between the secreted and membrane-bound PSG9 isoforms in PE. This bears resemblance to the imbalance between the soluble fms-like tyrosine kinase 1 (sFLT-1) and its cell membrane-bound isoform FLT-1 (VEGFR1) that has been implicated in PE pathogenesis [16]. Based on our data, it is tempting to speculate that while the membrane anchored PSG9 isoform modulates the function of the mature STBs, the soluble form promotes the signalling process for syncytialization as a ligand. This later model mirrors a ‘positive-feedback loop’, in which the soluble hCG synthesized in STBs binds to the LH chorionic gonadotropin receptor (LHCGR) and reinforces the cAMP signalling pathway [114, 115]. Our data also suggest that the secreted isoform may also influence the invasiveness of the extravillous trophoblasts. Shallow trophoblast invasion during placentation has been previously associated with PE [16–18].

From the LTR8B/PSG9 perspective, our data show that while PSG9 was primarily associated with extravillous trophoblast functions [59, 63, 116], LTR8B-driven PSG9 is predominantly expressed in syncytiotrophoblasts (STB), where it supports typical STB functions: PSG9 defines STB identity and possibly modulates the secretion of chorionic gonadotropin, the hCG hormone (more evidence is needed as regards this later point). Interestingly, the dysregulated LTR8B/PSG9 locus, which controls multinucleated cell differentiation, and several key pregnancy genes, including trophoblast-specific TFs, are likely to have an even greater impact on the pathological phenotype in addition to the potential direct effects of increased PSG9 gene products (e.g. on invasion) in EO-PE. Supporting this, in patient data the correlation of combined GATA3 and DLX5 dosage with PSG9 has a p-value < 0.01, whereas Fisher’s combined p-value test for the two individual effects yields p = 0.039, suggesting an interaction between GATA3 and DLX5 in regulating PSG9.

According to our analysis, both GATA3 and DLX5 contribute to the control of trophectoderm-specific regulatory networks of the early human embryo. Via these early developmental TFs, LTR8B/PSG9 may participate in evolutionary processes at the interface of embryogenesis and placentation. Dysregulation of DLX5 and GATA3 has been linked to EO-PE [38, 117]; however, the downstream genes and pathological phenotypes associated with their overexpression show partial but not complete overlap, reflecting the heterogeneity of PE. Notably, both DLX5 and GATA3 bind to the LTR8B/PSG9 platform, and PSG9 overexpression correlates more strongly with their combined dysregulation. Interestingly, DLX5 overexpression alone does not lead to the upregulation of LTR8B/PSG9 in the chromatin context, yet in patient datasets, PSG9 levels do increase with DLX5 expression, and PSG9 is reduced in DLX5-depleted, differentiated TSCs. A potential explanation is that DLX5 binding may need to precede recruitment of the pioneer transcription factor GATA3 at the LTR8B/PSG9 locus; in its absence in cells lines, DLX5 expression may be insufficient to induce PSG9. This aligns with our model that TF activity at the LTR8B/PSG9 locus is modulated by co-factor availability. Supporting this, in patient data the correlation of combined GATA3+ DLX5 dosage with PSG9 has a p value < 0.01, whereas Fisher’s combined p-values test for the two individual effects yields p = 0.039, suggesting an interaction between GATA3 and DLX5 in regulating PSG9.

### Why are ERVs so commonly associated with placental evolution?

Our evidence strengthens the link between ERVs and the evolution of placentation. While the Syncytin 1 and 2 genes are co-opted retroviral *env* genes that play an important role in human placentation [20, 28, 36, 55], the PSG9 locus provides a new example, demonstrating that ERV co-option events (both LTR8B and MER65) are central to the evolution of human pregnancy. The LTR8B/PSG9 locus represents a case in which an ERV-derived CRE becomes an essential player in a complex physiological process, such as the differentiation of multinucleated trophoblasts (*forskolin-stimulated cyclic AMP (cAMP) signalling*) and their placental function. Interestingly, our study shows that the LTR8B/PSG9 CRE and not the PSG9 gene per se is the essential regulator of the syncytial trophoblast differentiation process and thus a novel key player in human placentation.

One reason for this commonality may well be comparable to the “guns for hire” hypothesis [118] to explain why old ERVs are often recruited as suppressors of other invasive elements (e.g. in ref [119]). In this hypothesis the logic is, in part, that any and every suppressor is selectively favourable and that old ERVs are often predisposed to aid suppression by virtue of many of the features that once made them successful elements. In the context of placentation, rapid evolution possibly owing to maternal-foetal conflicts (as opposed to host parasite conflict) is comparable to the rapid evolution of suppressors of transposable elements. LTR8B’s involvement is then transparent: a successful ERV needs to recruit host TFs and the LTRs are the sites of that recruitment. Their random insertion next to some genes has utility for the foetal host, next to others not so much. The former are selectively favoured, the latter more likely to decay. Whether MER65 requires a polyadenylation signal as part of its host invasion is less transparent. The guns for hire hypothesis would predict that it may well have been important.

## Conclusions

Our study emphasises the evolutionary significance of endogenous retroviral elements (ERVs), specifically LTR8B and MER65, in regulating the pregnancy-specific glycoprotein (PSG) family, particularly PSG9, within the context of placental development and the pathology of pre-eclampsia (PE). The presence of LTR8B and MER65 elements within the PSG array has likely driven rapid evolutionary adaptations, enabling functional diversity among PSG isoforms. Through multi-omics and CRISPR/Cas9 analyses, we identified LTR8B as a trophoblast-specific CRE at the PSG9 locus, essential for driving PSG9 expression and syncytiotrophoblast differentiation. Of note, Based on chromatin accessibility, histone marks, transcription factor binding, and Hi-C data, we initially classified LTR8B as a candidate cis-regulatory element (cCRE). However, our CRISPR–Cas9 knockout experiments, which led to a complete loss of PSG9 transcription, provide direct functional validation that LTR8B acts as a bona fide cis-regulatory element (CRE) at the PSG locus. This highlights that, alongside the ERV envelope gene-derived syncytins, ERV regulatory elements have been integral to placental evolution. We further demonstrated that PSG9, unique among the PSG family, possesses both secreted and membrane-bound isoforms, a feature likely facilitated by MER65’s alternative polyadenylation signal, which distinguishes PSG9 from other family members and supports its secretory functions in the placenta.

Additional studies will be needed to clarify how PSG9 target genes mediate LTR8B’s influence on syncytiotrophoblast differentiation. Future research should also assess PSG9’s utility as an early-stage pregnancy biomarker and continue to explore the broader roles of ERVs in placental development and pathology.

Our findings suggest that LTR8B binding by GATA3/ DLX5 underpins PSG9’s specific regulation, further implicating it in the dysregulation associated with PE. Elevated PSG9 levels correlate with dysregulated GATA3 and DLX5 in early-onset PE (EO-PE), positioning PSG9 as a potential predictive biomarker. Moreover, we propose that the distinct regulatory network surrounding LTR8B/PSG9, involving chromatin interactions and specific transcription factor binding, is pivotal for syncytiotrophoblast identity and function, and its dysregulation may contribute to PE pathology. Additional studies will be needed to clarify how PSG9 target genes mediate LTR8B’s influence on syncytiotrophoblast differentiation. Future research should also assess PSG9’s utility as an early-stage pregnancy biomarker and continue to explore ERV roles in placental development and pathology.

## Acknowledgments

We thank all members from Izsvak lab for their support and suggestions in the project.

## Authors’ contributions

M.S., Z.I., J.Z., and conceived the idea and designed the study. Y.Q. and J.Z. performed most of the experiments. F. H., M.G., X. K., R.Z., R.A., and K.S. assisted with the experiments. F.H handled cohorts and supervised/designed expression studies and primary cell isolation. M.K.K. has performed the CNV analysis. M.G. performed immunohistochemistry. G.D. assisted with the genome wide sequencing. S.M.B established the PSG9-sepcific ELISA. M.S., performed all the high throughput data analysis except the CNV and alternative transcript analysis of PSG9. A.P. performed RNA-seq, CNV and alternative transcript of PSG9 analysis. L.D.H. assisted some statistical analysis. Z.I., M.S., Y.Q. and L.D.H. wrote the manuscript with input from all the other authors. All authors have read and approved the final manuscript.

## Funding

This work was supported by the following funding sources: Y.Q was supported by the China Scholarship Council (NO. 201504910684) and Guangdong Basic and Applied Basic Research Foundation (2025A1515012066). M.G. was supported by the Austrian Science Fund (FWF): 10.55776/I3304 and 10.55776/I6907.

## Data Availability

All original code has been deposited at Code availability Homemade R script used in this study is publicly available on GitHub (https://github.com/Manu-1512) and v1.0.0 amitpande74/PSG9-Isoform-Transmembrane-Topology-Prediction-using-ProtBERT: PSG9 ProtBERT Analysis v1.0; DOI: 10.5281/zenodo.17084782; PSG9 ProtBERT Analysis v1.0

The public datasets analysed in this study are listed in Additional File 1.

## Declarations

For the ethical approval see Additional File 3: Patients and clinical data

## Consent for publication

Not applicable.

## Competing interests

The authors declare no competing interests.

## Notes

### Competing Interest Statement

The authors have declared no competing interest.

