## Supplementary Materials for "Endogenous retroviral elements LTR8B and MER65 regulate the PSG9 locus that promotes trophoblast syncytialization: Insights into placental evolution and pre-eclampsia pathology"

**Additional File 3: Patients and clinical data**

PE was defined as women with the new onset of hypertension (systolic blood pressure of at least 140 mmHg or more and/or diastolic blood pressure of at least 90 mmHg) after 20-week gestation, accompanied by proteinuria (≥ 0.3 g protein in a 24-hour urine specimen) [101]. Pregnant women with multiple gestations, smoking history or chronic diseases (e.g., diabetes, thyroid disease, kidney dysfunction, autologous Immune diseases) were excluded from the current study. Early-onset PE is defined as delivery < 34 gestational weeks. Late-onset PE is defined as delivery ≥ 34 gestational weeks. BMI: body mass index. The following cohorts were used in this study:

**Oslo cohort:** The placental samples are comprising of patient samples from bio-bank collection at Oslo University Hospital, Norway. Samples from the Oslo cohort were analysed in [102, 103] The Oslo cohort consists of placental tissues collected during elective Cesarean sections in 47 PE patients and 27 controls with normotensive and uncomplicated pregnancies. The PE group was further divided into early onset (EO) PE (delivery < 34 gestational weeks, n = 24) and late-onset (LO) PE (delivery ≥ 34 gestational weeks, n = 23 samples. The uncomplicated pregnancy group consisted of healthy, normotensive women undergoing elective Cesarean section due to breech presentation or other reasons. The Oslo Pregnancy Biobank samples stem from an ongoing recruitment of pregnant patients. The study (and the present research collaboration) is approved by the Regional Committee of Medical Research Ethics South Eastern Norway (ref 2013/2092 and 529-02162). Informed written consent was obtained from each participant.

| Oslo Characteristics at delivery | Controls (n=27) | PE Early-onset (n=24) | Statistics |
|---|---|---|---|
| **Maternal age (Years)** | 31.2 ± 4.2 | 31.6 ± 5.6 | P=0.63 |
| **BMI ($kg/m^2$)** | 28.6± 3.4 | 31.5 ± 5.2 | **P ≤ 0.01 |
| **Gestational week** | 39.0 ± 0.9 | 33.7 ± 3.7 | ***P ≤ 0.001 |
| **Systolic BP (mm Hg)** | 119 ± 11.6 | 165 ± 16.2 | ***P ≤ 0.001 |
| **Diastolic BP (mm Hg)** | 72.1 ± 10.9 | 101.1 ± 6.7 | ***P ≤ 0.001 |

| Baby weight (g) | 3492± 402 | 2170 ± 1005 | ***P ≤ 0.001 |
|---|---|---|---|

**Charité-Aachen cohort:**

Preeclamptic and IUGR term placenta samples and gestational age matched controls were collected in the University Hospital of the RWTH Aachen, Germany (as described above) and at Charité – Universitätsmedizin Berlin. Sampling was approved by the local ethical committees (Aachen: EK 148/07; Berlin: EA2/132/12) and informed consent was obtained from each participating woman. Clinical characteristics of the PE, IUGR and control cases from the study population are listed in the table.

| Charite/Aachen Characteristics at delivery | Control early (n=36) | Control late (n=65) | IUGR early (n=28) | IUGR late (n=13) | PE Early onset (n=14) | PE late onset (n=23) |
|---|---|---|---|---|---|---|
| **Maternal age (Y)** | 31.4 ± 5.7 | 32.5 ± 5.6 | 27.7 ± 6.4 | 27.6 ± 5.8* | 33.0 ± 5.3 | 31.2 ± 6.2 |
| **BMI (kg/m²)** | 25.0 ± 6.1 | 23.6 ± 4.6 | 24.0 ± 4.8 | 26.2 ± 7.0 | 26.0 ± 5.1 | 27. ± 7.0* |
| **Gestational days** | 203.3 ± 22.1 | 264.7 ± 13.0 | 208.6 ± 19.0 | 264.3 ± 11.8 | 206.1 ± 21.3 | 254.5 ± 12.0 |
| **Systolic BP (mm Hg)** | 115 ± 9.7 | 116.6 ± 10.8 | 122.9 ± 13.7*** | 104.5 ± 15.5* | 152.6 ± 12.4*** | 153.5 ± 14.5*** |
| **Diastolic BP (mm Hg)** | 65.0 ± 8.0 | 69.7 ± 8.1 | 70.7 ± 11.7*** | 65.2 ± 10.3 | 95.9 ± 8.7*** | 94.8 ± 8.8*** |

Data are presented as mean ± standard deviation. *P ≤ 0.05, ***P ≤ 0.001 vs. control.

**Manchester Antenatal Vascular Service Cohort**

Maternal plasma samples included in this study were collected from 24th - 28th weeks of gestation from a high-risk cohort in the United Kingdom, the Manchester Antenatal Vascular Service (The MAViS clinic). The clinical characteristics have been reported in detail previously. Informed written consent was collected from each participant. The study was approved by the NRES (National Research Ethics Service) Committee North West 11/NW/0426. The inclusion criteria for women in the MAViS study were: (1) chronic hypertension BP ≥140/90 at ≤20 weeks; (2) chronic hypertension requiring antihypertensive treatment ≤20 weeks; (3) pre-gestational diabetes mellitus with evidence of vascular complications (hypertension, nephropathy); (4) history of ischaemic heart disease; and (5) previous early onset preeclampsia.

**Hungary cohort**

Serum samples were obtained from women with uneventful pregnancies (healthy controls) and women with preeclampsia at the First Department of Obstetrics and Gynaecology, Semmelweis University, Budapest, Hungary. The study was approved by the Regional and Institutional Committee of Science and Research Ethics at Semmelweis University (IRB No. 188/2008), and written informed consent was obtained from all participants. The research was conducted in accordance with the principles of the Declaration of Helsinki. All participants were Caucasian and resided in the same geographic region of Hungary. The inclusion criteria required women to be at least 18 years old and to have a singleton pregnancy. Exclusion criteria included multifetal gestation, pre-existing chronic hypertension, diabetes mellitus, autoimmune diseases, angiopathies, renal disorders, maternal or fetal infections, or fetal congenital anomalies. Preeclampsia was diagnosed according to the following criteria: systolic blood pressure ≥140 mmHg or diastolic blood pressure ≥90 mmHg on at least two occasions, separated by ≥6 hours, after 20 weeks of gestation in a previously normotensive woman, in conjunction with proteinuria (≥0.3 g/24 hours or ≥1+ on a urine dipstick test, excluding urinary tract infections). Clinical characteristics of the study groups are summarized in the Table below.

| Hungary Cohort Characteristics | Controls (n=92) | PE (n=67) | Statistics |
|---|---|---|---|
| **Maternal age (Years)** | 31.4 ± 3.9 | 31.2 ± 4.5 | ns |
| **BMI (kg/m$^2$)** | 27.4± 4.3 | 29.7 ± 3.4 | *P ≤ 0.05 |
| **Gestational week** | 36.0 ± 0.8 | 35.5 ± 0.7 | ns |
| **Systolic BP (mm Hg)** | 120 ± 4.5 | 150 ± 10.1 | ***P ≤ 0.001 |
| **Diastolic BP (mm Hg)** | 70.2 ± 9.3 | 100.9 ± 8.6 | ***P ≤ 0.001 |
| **Baby weight (g)** | 3415 ± 480 | 2575 ± 1210 | ***P ≤ 0.001 |
