## Supplementary Figures for "Endogenous retroviral elements LTR8B and MER65 regulate the PSG9 locus that promotes trophoblast syncytialization: Insights into placental evolution and pre-eclampsia pathology"

**Fig. S1 Conservation LTR8B and MER65-int element in primates.** UCSC browser snapshot showing the comparative genomics (GRCh38/hg38) of the PSG9 locus. The LTR8B and MER65-int are specifically present in anthropoid primates (light blue).

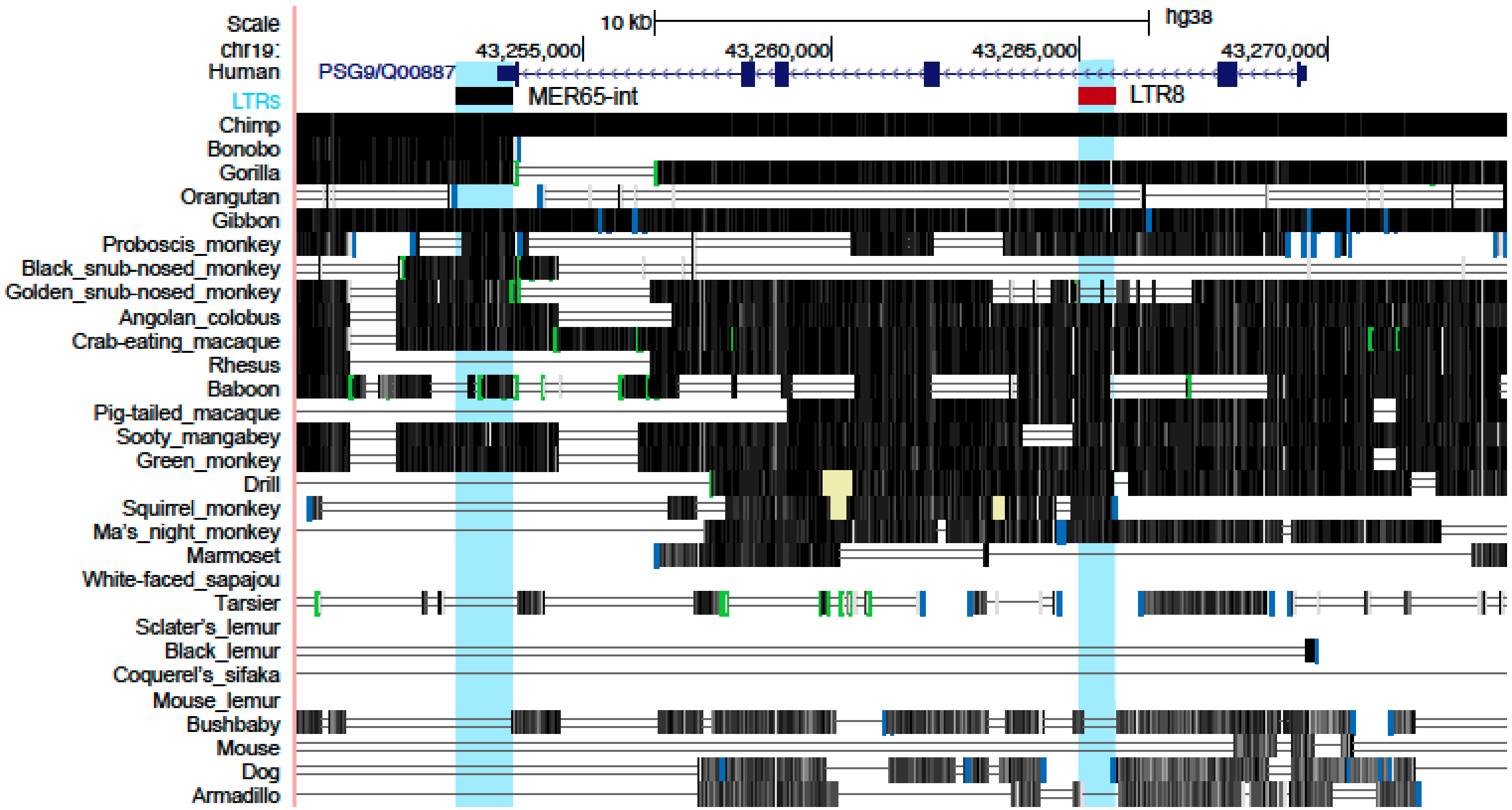

**Fig. S2 Specificity of the PSG9 antibody.** Specificity studies of PSG9 antibodies using Western blotting. (Left panel) Purified recombinant PSG proteins [58] were loaded on the gels 2 µg /lane: PSG1 V5His, PSG3 V5His, PSG4 V5His, PSG5 V5His, PSG6 V5His, PSG7 V5His, PSG8 V5 His, PSG9 V5His and PSG11V5 His. Primary Ab Novus rabbit polyclonal NBP-2 19979 at a 1:1,000 dilution, followed by goat anti-rabbit HRP conjugated Ab. Note that this Ab is advertised as PSG9-specific but it is not [43]. (Right panel) 1.5 µg / lane of recombinant PSG proteins PSG1 V5His, PSG2 V5His, PSG4 V5His, PSG5 V5His, PSG6 V5His, PSG7 V5His, PSG8 V5 His, PSG9 V5His and 20 µg lysates of BeWo, SHGPL-4 cells. Primary Ab was rabbit polyclonal anti-PSG9 (ab64425) 1:1000, followed by anti-rabbit-HRP 1:5000.

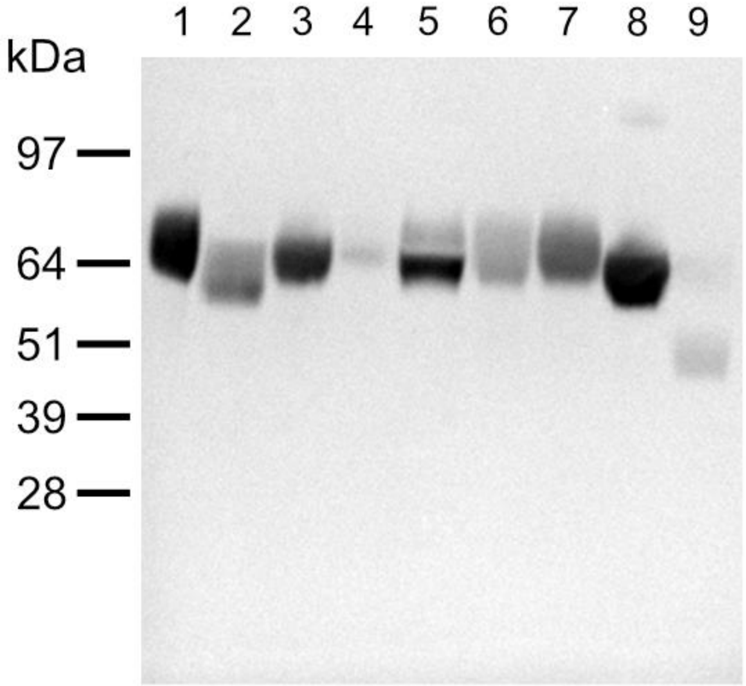


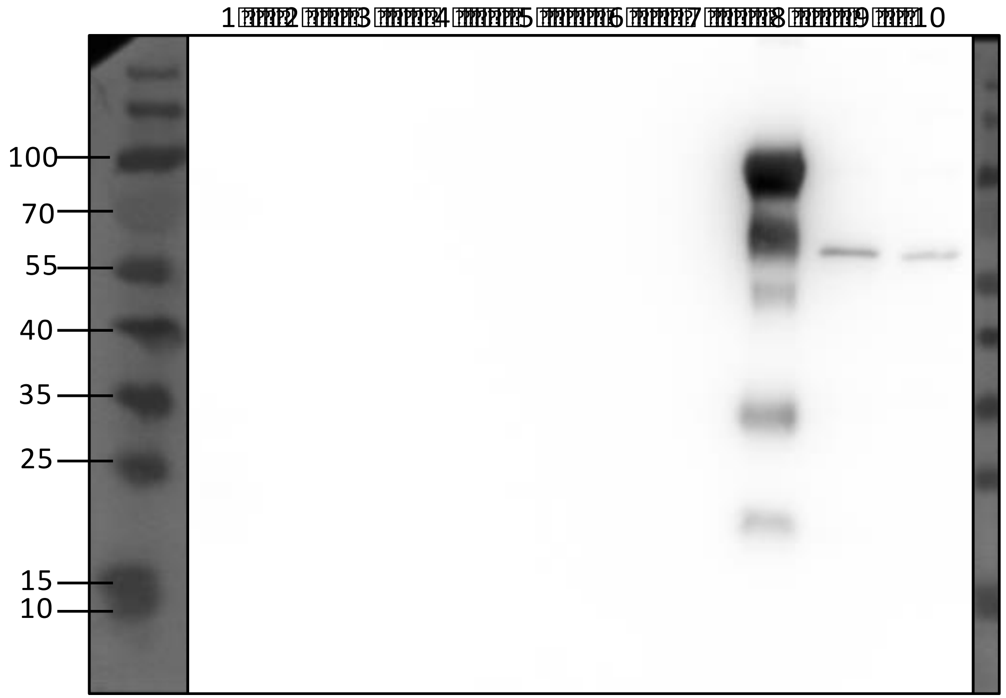


Fig S2

**Fig. S3** Schematic of the luciferase reporter assay to analyse TE sequences for enhancer activity.

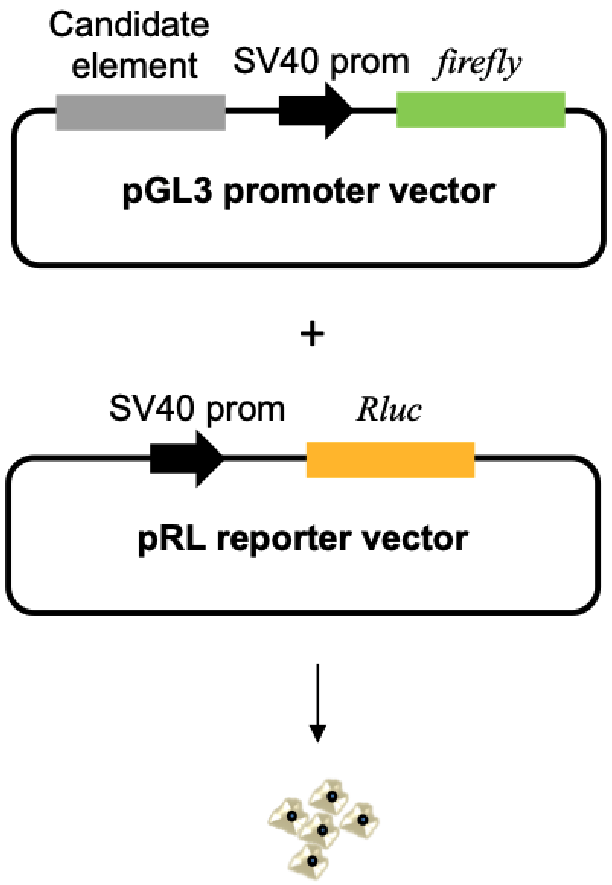

**Fig. S4 Characterisation of the LTR8B/PSG9 locus.** Chromatin interaction profiling of the *PSG9* gene region. The upper panel shows the arc-diagram of the identified interactions with the *P*-value < -log10 (2), interaction range = 1Mb. The lower panel shows the genes in the interaction range. (Bottom) Genome browser snapshot showing the chromatin features of the genomic region overlapping with the LTR8B in the PSG9 2nd intron in human PSC-derived trophoblasts. One-to-all interaction frequency. Blue bar graph represents bias-removed chromatin interaction frequency, and magenta dots represent distance-normalized interaction frequency. Arc-diagram shows the identified interactions with the distance normalized interaction frequency >2, interaction range = 2Mb. (Bottom panel) shows the genes among the interaction range. The PC Hi-C shows the interaction of *PSG9* promoter/enhancer with *PSG4* promoter and other genomic regions (GRCh37/hg19). The arc-diagram shows the identified interactions with the *P*-value < -log10 (2), interaction range = 1Mb.

A

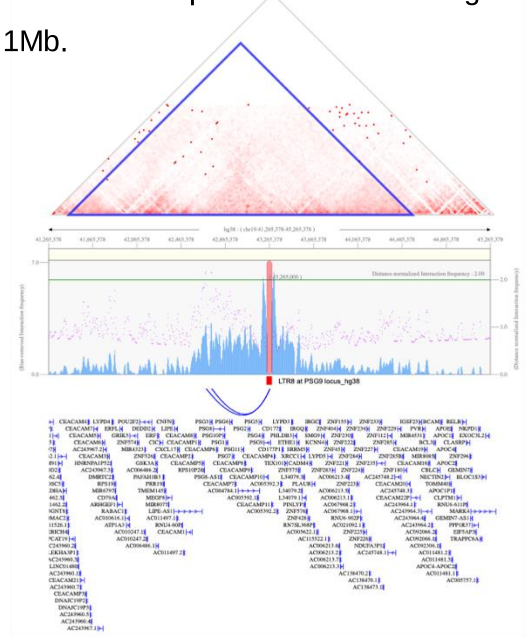


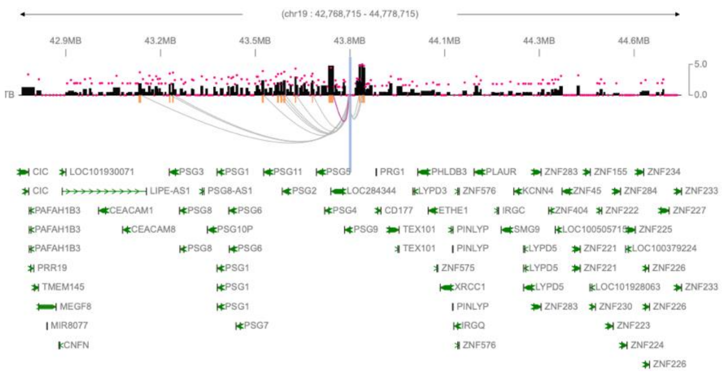

**Fig. S5 Knocking out LTR8B at the PSG9 locus (KO). A (Upper panel)** The CRISPR/Cas9-mediated deletion of LTR8B/PSG9 (990bp) using the guide RNA pair, shRNA1 and sgRNA3. Confirmed by genotyping and Sanger sequencing. **(Lower panel)** The CRISPR/Cas9-mediated deletion of LTR8B/PSG9 (690bp) using the guide RNA pair, shRNA2 and sgRNA3. Confirmed by genotyping and Sanger sequencing (Lower panel). **B** Downregulation of selected STB genes upon KO-LTR8B. **C** Downregulation of selected genes, involved in STB differentiation. **D** Deletion of LTR8B/PSG9 results in increased levels of CDX2 and TEAD4.

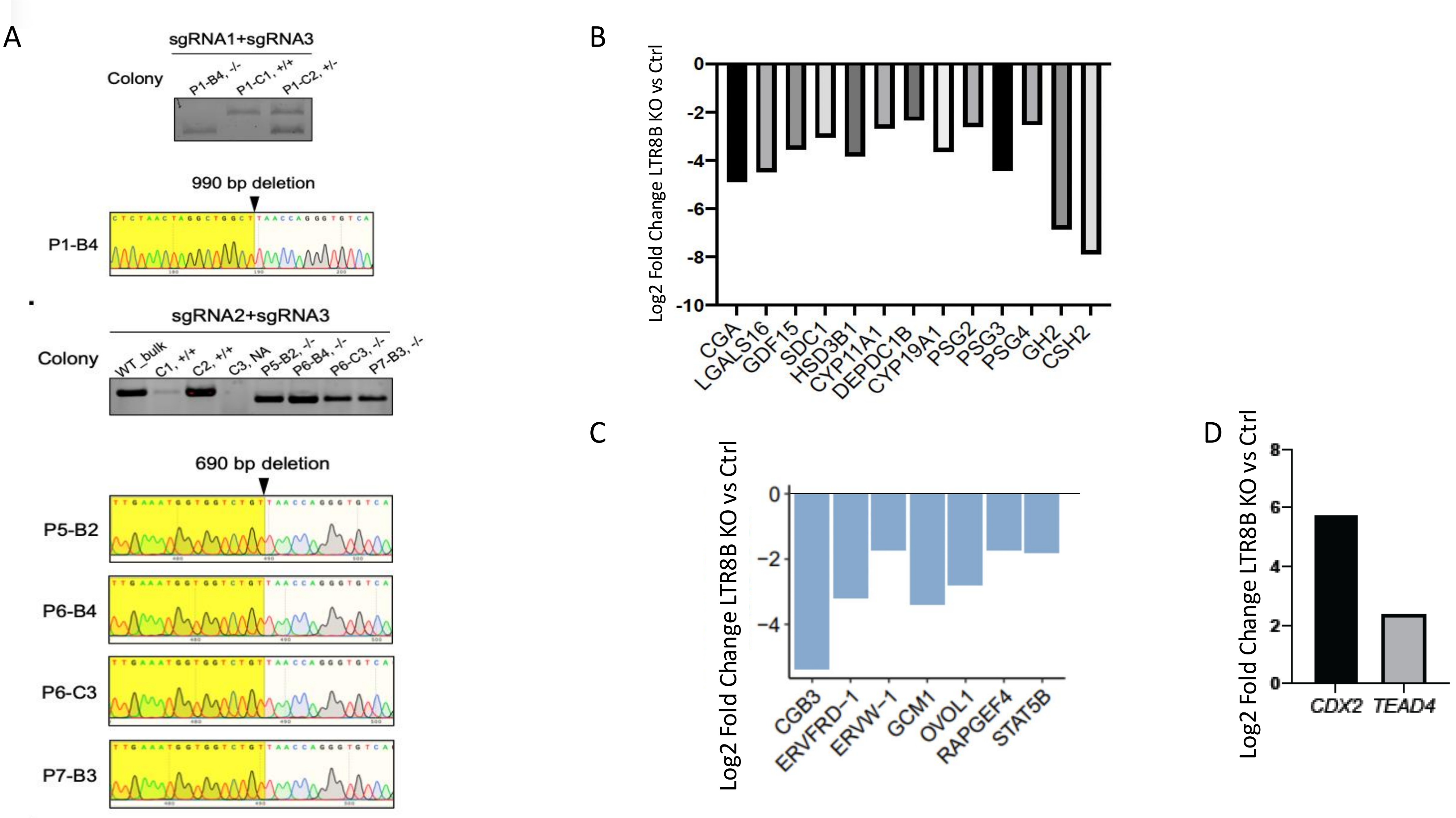


Fig S5

**Fig. S6 Impact of the ectopic expression of trophoblast specific TFs on PSG9/LTR8B. A (Top Panel)** Validation of the ectopic over-expression (OE) of HA-tagged TFs (HA-GATA3 and HA-TFAP2A) by Western blotting (representative image of three independent replicates). **(Bottom Panel)** RT-qPCR analysis on the overdosing effect of GATA3 and TFAP2A TFs on the reporter gene expression in STBs. N = 8 technical replicates, mean $\pm$ SD. Ordinary one-way ANOVA followed by multiple comparation. Control vs. GATA3 or TFAP2A, **** $P < 0.0001$. **B** Validation of the ectopic over-expression (OE) of HA-tagged TFs (HA-GATA2, HA-GATA3, HA-TFAP2A, HA-TFAP2C). **C** RT-qPCR analysis on the overdosing effect of various TFs on the expression of PSG1 in STBs. N = 3 independent replicates, mean $\pm$ SD. One-way ANOVA followed by multiple comparation. The measured differences were not significant (level $P < 0.05$).

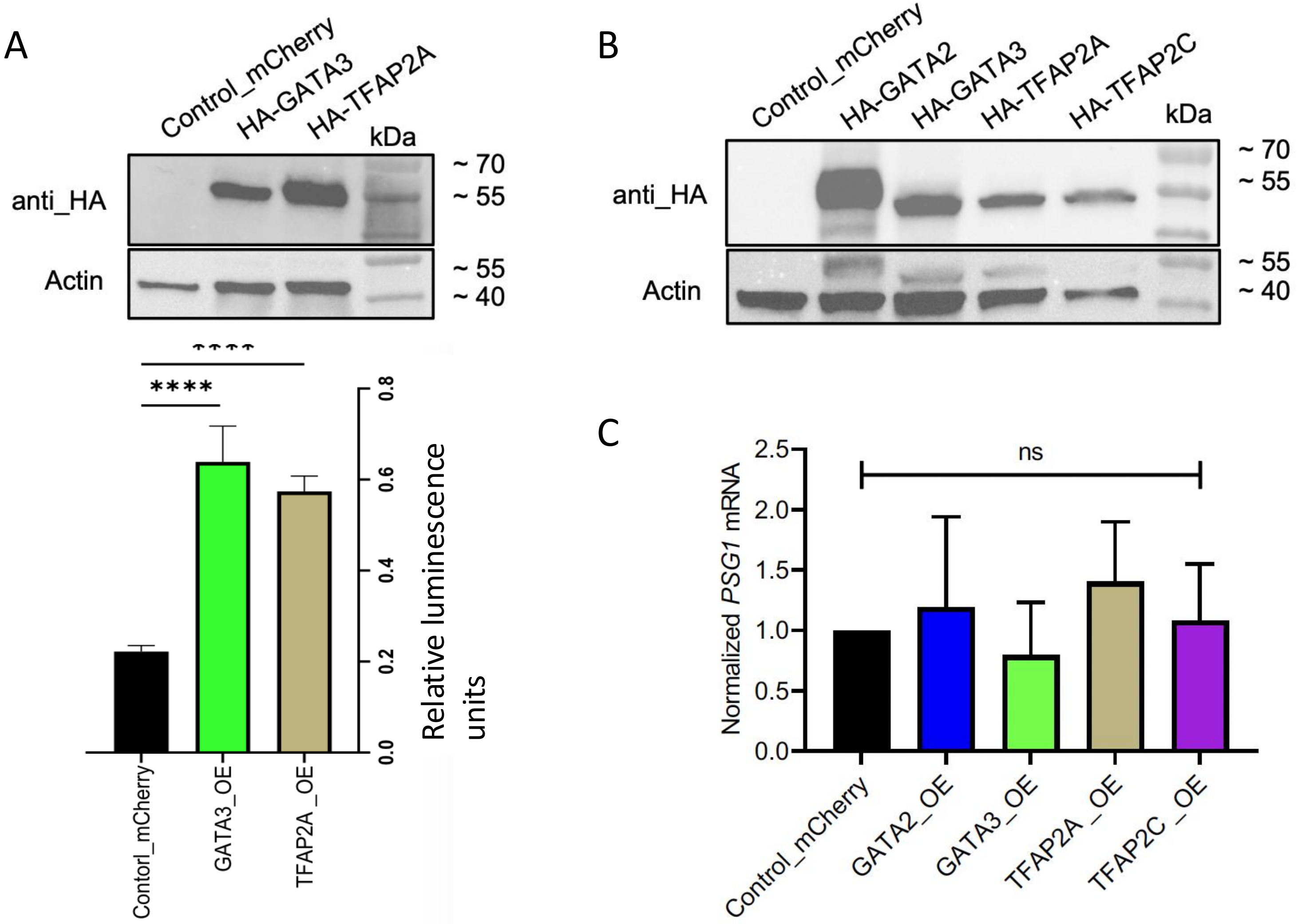


Fig S6

**Fig. S7 Annotated PSG9 isoforms.** The consensus polyA sequence is embedded in MER65-int element.

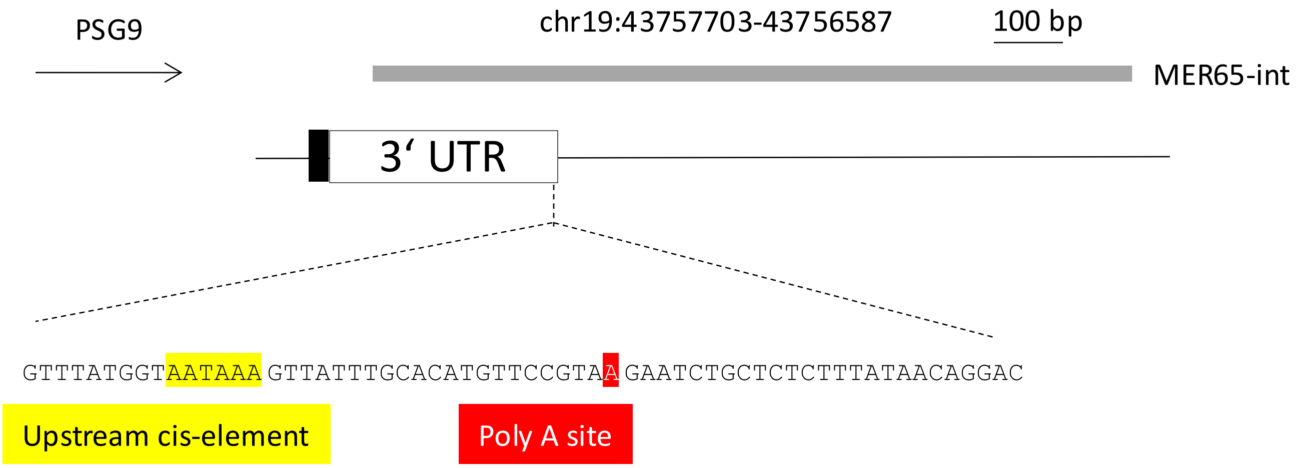

**Fig. S8 MER65 defines subcellular localization of PSG9. A** Schematic of the luciferase reporter assay to validate the polyA signal embedded in MER65-int. **B** Hydrophobicity analysis of the PSG9-202 and PSG9-201 isoforms. Note: **C** Transmembrane prediction of the PSG9-202 and PSG9-201 isoforms (TMHMM probabilities).

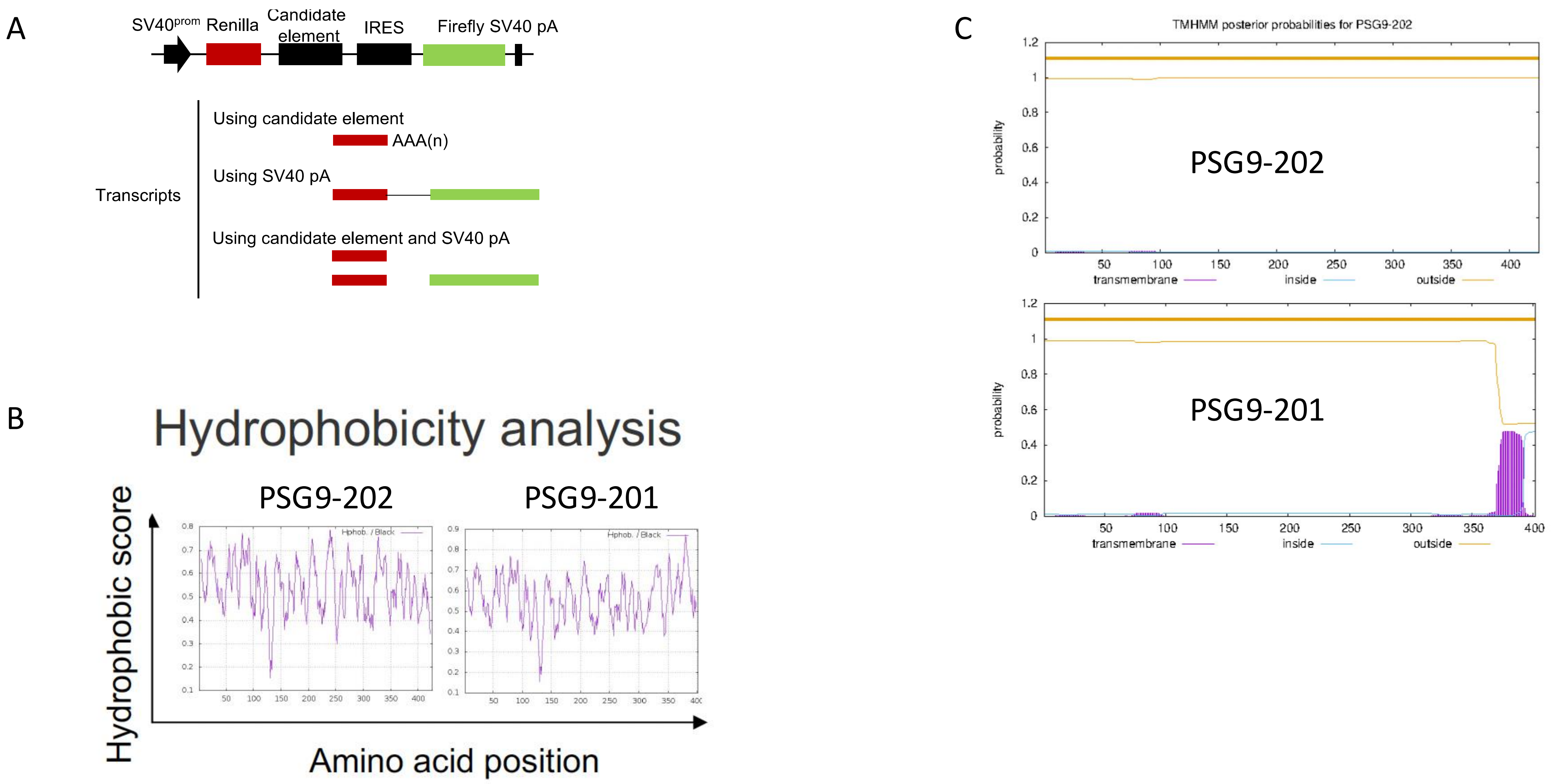

**Fig. S9 Characterisation of the membrane bound and secreted PSG9 isoforms. A** Pairwise comparison of PSG9-201 and PSG9-202 amino acid sequences. Note that PSG9-201 lacks the A2 domain but contains a predicted transmembrane region. **B** The strategy of marking PSG9-201 and PSG9-202 protein variants with HA and FLAG tags, respectively, and stably coexpress them in BeWo cells with the help of the *Sleeping Beauty* transposon system.

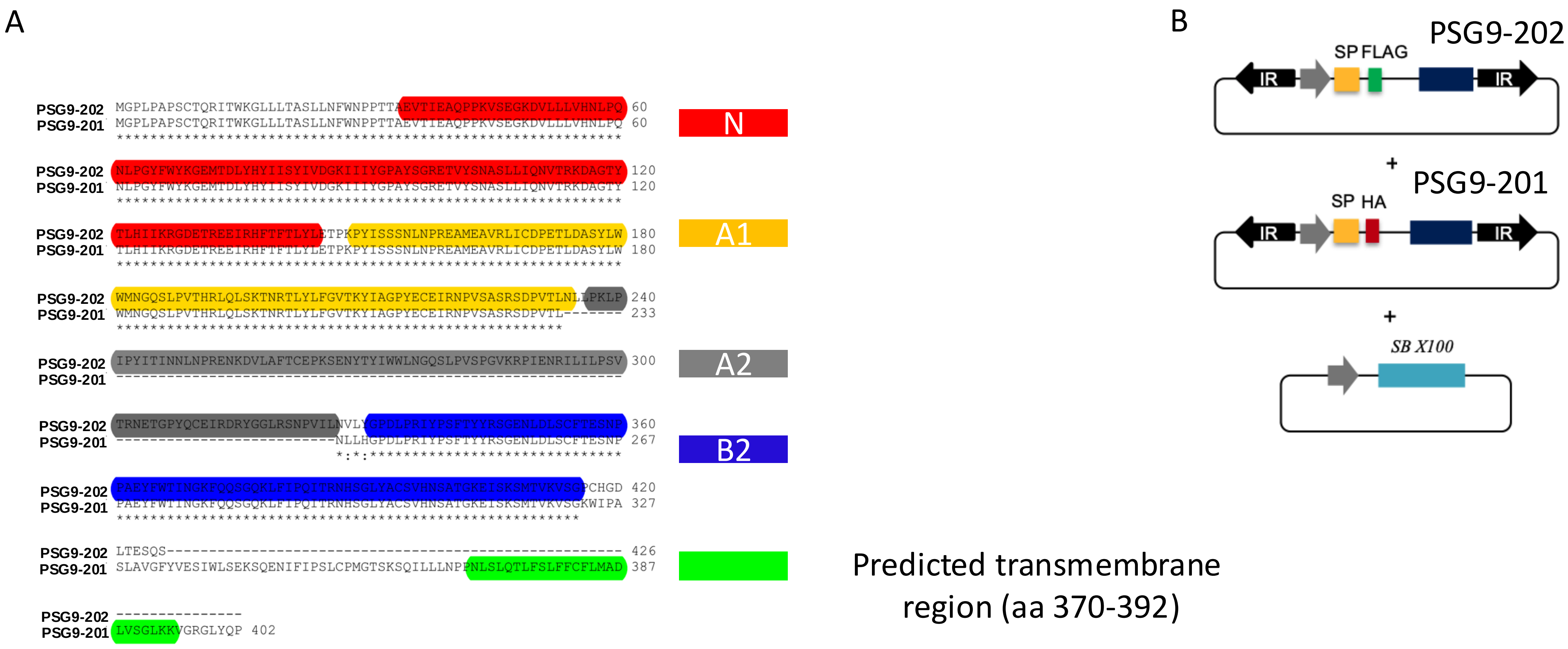
